# Tubulin autoregulation factors SCAPER and TTC5 recruit γ-tubulin to non-centrosomal MTOCs for neuronal microtubule nucleation and axon regeneration

**DOI:** 10.1101/2025.10.13.682208

**Authors:** Yi Xie, Alessandro Comincini, Chengye Feng, Carrie Ann Davison, Marc Hammarlund, Francesca Bartolini, Shaul Yogev

## Abstract

Neuronal function and survival depend on highly stereotyped non-centrosomal microtubule (MT) arrays. How these arrays form remains poorly understood. Here we identified a role for SCAPER and TTC5, two factors previously implicated in *tubulin* mRNA autoregulation, in controlling neuronal MT content through γ-tubulin-dependent nucleation. In *C. elegans* neurons, loss of *scpr-1, ttc-5,* or both reduced MT numbers to a similar degree as depletion of γ-tubulin, the main MT nucleator. Using conditional single-cell degradation alleles and endogenous tagging, we find that γ-tubulin nucleates MTs in the neuronal cell body from endosomal puncta, and that *scpr-1* and *ttc-5* are required to recruit γ-tubulin to these structures. SCPR-1 is also instructive, as its overexpression drastically increases γ-tubulin levels and enhances MT density. We propose that these mechanisms are conserved since human SCAPER rescues *C. elegans* mutants, and SCAPER knockdown in rat hippocampal neurons reduces both γ-tubulin clustering at presynaptic sites and activity-dependent synaptic MT nucleation. Finally, while *scpr-1, ttc-5,* and *γ-tubulin* are not required for developmental axon elongation, they are essential for regeneration, where SCPR-1 directs γ-tubulin to the growth cone to facilitate regrowth following injury. These findings reveal mechanisms governing neuronal cytoskeleton assembly and function, and suggest potential crosstalk between tubulin autoregulation and microtubule nucleation.

## Introduction

Neuronal viability and function rely on a specialized MT cytoskeleton. In axons and dendrites, tiled arrays of non-centrosomal MT polymers support long-range transport and are essential for growth or regeneration ^1–4^. The architecture of these arrays – polymer numbers, length, and spatial distribution – is highly stereotyped within neuronal subtypes ^5,6^. For example, *C. elegans* motor neurons and mouse Purkinje parallel fibers contain sparse MT arrays, whereas worm touch receptor neuron axons and cultured rodent hippocampal axons and dendrites harbor dense MT networks ^7–10^. While this stereotypy suggests that MT content – defined here as the total number of MT polymers in a cell – is tightly regulated, the underlying mechanisms remain poorly understood.

MT nucleation initiates polymer assembly by overcoming the kinetic barrier to tubulin polymerization, and its regulation is likely key for determining cellular MT content ^11–13^. The major MT nucleator, γ-tubulin ring complex (γ-TuRC), consists of γ-tubulin and associated proteins that together provide the template for nascent MT assembly ^14–19^. γ-TuRC is an inefficient nucleator *in vitro*, and in cells it needs to be recruited and concentrated at specialized locations such as the centrosome, where it promotes the formation of radial MT arrays ^20,21^. In addition to the centrosome, neuronal γ-TuRC also localizes to somatic Golgi, Golgi outposts and endosomes ^22–26^ that act as non-centrosomal MTOCs. Centrosomal recruitment occurs early in development, mediated by factors such as NEDD1 and CDK5RAP2, and is important for neuronal migration ^27–29^. Non-centrosomal MT-organizing centers establish mature MT arrays in axons and dendrites and play important roles in controlling MT polarity, dynamicity, and functions such as supporting cargo transport and synaptic function ^30–34^. However, the mechanisms controlling the assembly of non-centrosomal MTOCs in neurons remain poorly understood.

In addition to nucleation, the regulation of tubulin homeostasis could control cellular MT content. One pathway that regulates tubulin homeostasis is autoregulation: in response to experimental elevation of soluble tubulin levels, the autoregulation factors TTC5 and SCAPER are recruited to *tubulin-*translating ribosomes and destabilize the bound *tubulin mRNA* through recruitment of the CCR4-NOT complex ^35–38^. While tubulin autoregulation is understood at a mechanistic level, its physiological role remains unclear since it is unknown how it is endogenously triggered in cells. Interestingly, the phenotypic spectrum of patients harboring mutations in both TTC5 and SCAPER suggests an important role for these proteins in neurodevelopment ^39–41^. However, precisely what that role is and whether it involves autoregulation and control of neuronal MT architecture is unknown.

Here we found that the tubulin autoregulation factors TTC-5 and SCPR-1 play an important role in controlling the neuronal MT network architecture by promoting γ-tubulin-mediated nucleation. In *C. elegans,* loss of *scpr-1* leads to a 30% reduction in the number of polymers, and in rat hippocampal neurons SCAPER is required for activity-dependent nucleation of synaptic microtubules. *scpr-1, ttc-5* and *γ-tubulin* function in the same genetic pathway and colocalize on endosomal compartments, where they promote MT nucleation. TTC-5 and SCPR-1 mediate the recruitment of γ-tubulin to these structures and SCPR-1 can also instruct excessive γ-tubulin recruitment and MT nucleation. Human SCAPER rescues *scpr-1* mutants in *C. elegans,* and rat SCAPER is required for the synaptic enrichment of γ-tubulin, suggesting that it functions similarly to *C. elegans* SCPR-1. Finally, γ-tubulin recruitment by SCPR-1 also occurs in growth cones of regenerating axons, where it is important for regrowth following injury. These findings provide insight into regulatory mechanisms that control neuronal MT architecture and function.

## Results

### *scpr-1* is required for neuronal microtubule organization

Microtubules (MTs) are crucial for neuronal function and survival, but how the density of the neuronal MT array is regulated remains largely unclear. To uncover regulators of neuronal MT organization, we adapted the *C. elegans* cholinergic motor neuron DA9 as a system that is amenable to genetic screens. DA9 is located in the preanal ganglion, from where it grows an anterior dendrite and a posterior axon that traverses to the dorsal cord before turning anteriorly and extending ∼500 μm by the L4 larval stage (**Figure 1A**). We visualized MTs using transgenes for the minus-end-associated protein PTRN-1/CAMSAP and TBA-1/α-tubulin (**Figure 1B**), which we have previously validated and shown to be reliable for estimating MT numbers, distribution, and average length ^42^. MTs in the DA9 dendrite and dorsal synaptic region were regularly distributed, with minus-ends spaced ∼1.4 μm apart, had an average length of ∼ 5 μm, and on average of 4-6 polymers are present per cross section, in agreement with previous EM reconstructions (**Figure 1I-L, Figure S1I-K**) ^7,42^. Similar polymer densities are observed in neurons from invertebrates to mammals ^43–46^.

**Figure 1.**
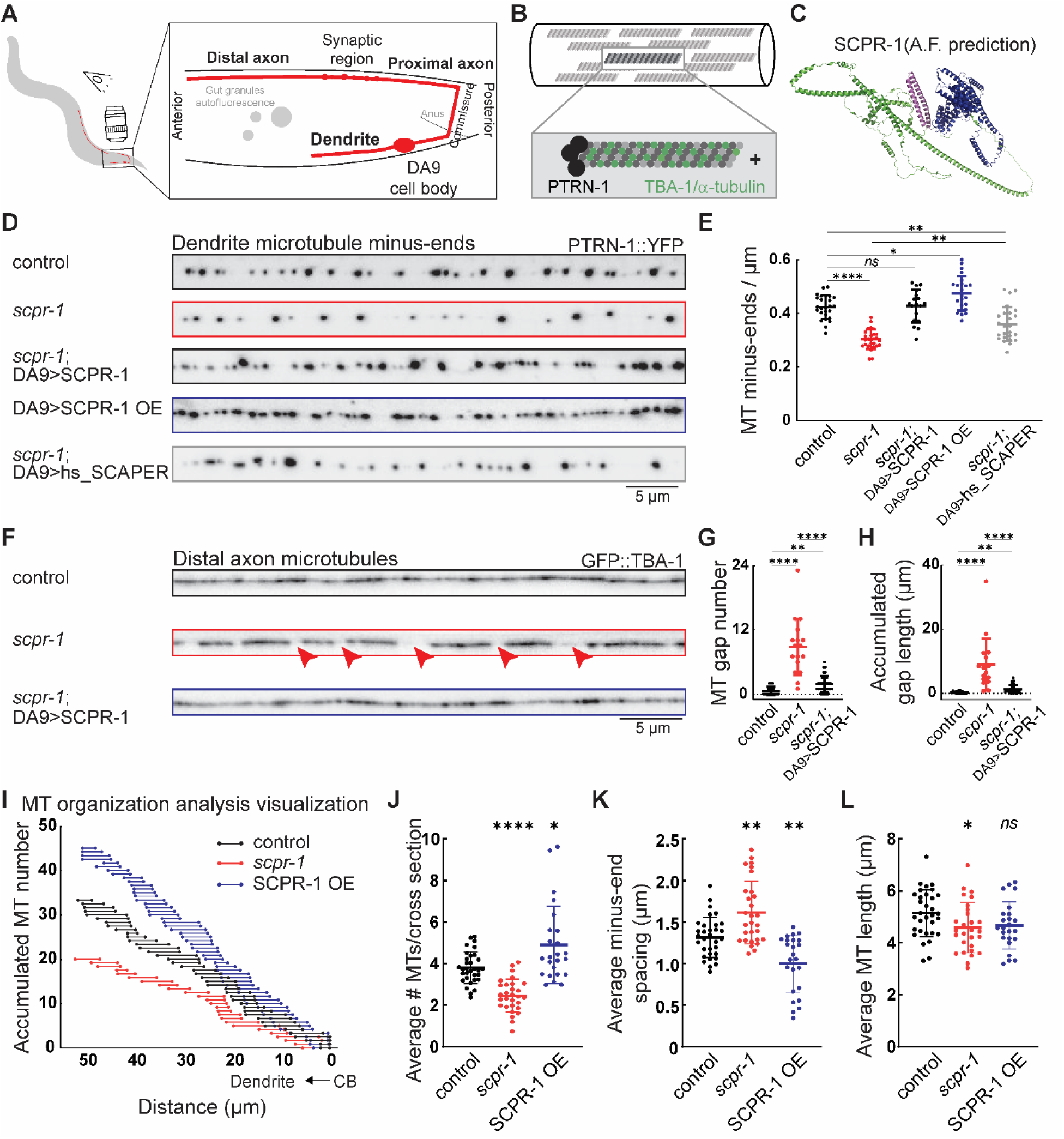
*scpr-1* is required for neuronal microtubule organization. (A) Schematic of C. elegans motor neuron DA9. (B) Schematic of microtubule arrays in DA9 neuron processes. MTs were labeled and visualized by MT minus-end protein PTRN-1 and tubulin subunit TBA-1. (C) AlphaFold structure prediction of C. elegans SCPR-1 (AF-G5EGE8-F1-model_v4). Magenta: N-terminus SCAPER domain. Green: α-helices domain. Blue: C-terminus domain. (D) Representative images of dendrite MT minus-end visualized by YFP-tagged PTRN-1 in control, *scpr-1*, *scpr-1* with DA9 SCPR-1 cDNA rescue, DA9 SCPR-1 overexpression, and *scpr-1* mutant with human SCAPER cDNA rescue, respectively. Scale bar = 5 μm. (E) Quantification of MT minus-end number density in DA9 dendrite in control (N=26), *scpr-1* (N=27), *scpr-1*; DA9>SCPR-1 (N=18), DA9>SCPR-1 OE (N=23), and *scpr-1*; DA9>hs_SCAPER (N=25). Brown-Forsythe and Welch ANOVA tests were applied. * (between control vs DA9>SCPR-1 OE): p=0.0137. **: p<0.005. ****: p<0.0001. *ns*: not significant. (F) Representative images of distal axon MT visualized by GFP-tagged MT subunit TBA-1 in control, *scpr-1*, and *scpr-1* with DA9 SCPR-1 cDNA rescue. Red arrow heads indicate gaps between MT polymers due to reduction of MT content. Scale bar = 5 μm. (G and H) MT gap number and accumulated gap length in control (N=18), *scpr-1* (N=17), and *scpr-1*; DA9>SCPR-1 (N=35). Kruskal-Wallis tests were applied. **: p<0.005, ****: p<0.0001. (I) Schematic of MT organization visualization from representative MT-Quant for control, *scpr-1*, and SCPR-1 OE. Each MT was illustrated by a single bar and aligned along with cell body to dendrite orientation horizontally in x-axis and accumulated MT number vertically in y-axis. (J-L) MT-Quant measurements and simulations of average number of MT per cross section, average MT minus-end spacing, and average MT length in DA9 dendrite for control (N=32), *scpr-1* (N=28), and SCPR-1 OE (N=24). Brown-Forsythe and Welch ANOVA tests were applied. * (SCPR-1 OE vs control in J): p=0.0304. * (*scpr-1* vs control in L): p=0.0231. **: p<0.005, ****: p<0.0001. *ns*: not significant.

From an unbiased genetic screen, we isolated *shy5*, a mutant showing a 30% reduction in the number of PTRN-1::YFP puncta (**Figure S1A-C**), suggestive of a reduced number of MT polymers. The *Shy5* phenotype is observed in both axons and dendrites and is shown for convenience in the dendrite in subsequent figure panels. We mapped *shy5* using genetic approaches and whole genome sequencing to a premature stop codon in a previously uncharacterized gene in *C. elegans,* which bears high sequence and structural homology to human SCAPER and was therefore renamed *scpr-1* (**Figure 1C**).

To study SCPR-1 function, we used CRISPR to delete exons 3-7 which contain a highly conserved SCAPER_N-terminal motif and confirmed by qRT-PCR that expression of the remaining sequence is strongly reduced (**Figure S1D**). *scpr-1* deletion phenocopied the reduction of PTRN-1::YFP in *shy5,* confirming the genetic mapping results (**Figure 1D-E**, **S1A-C**). Loss of *scpr-1* also reduced the number of PTRN-1 puncta in the Posterior Lateral Microtubule Neuron (PLM), a touch receptor neuron that harbors a dense MT array ^7^, indicating that SCPR-1 function is not restricted to DA9 (**Fig S1E-G**). Cell-specific expression of SCPR-1 cDNA in DA9 restored PTRN-1::YFP density in *scpr-1* mutants to control levels, indicating that SCPR-1 functions cell-autonomously (**Figure 1D-E**). Expression of human SCAPER in *scpr-1* mutants also increased PTRN-1 density in *scpr-1* mutants, suggesting that SCPR-1, is functionally conserved (**Figure 1D-E**). Last, overexpression of SCPR-1 in DA9 led to a 12.3% increase in the number of PTRN-1 puncta above control levels (**Figure 1D-E**). Together, these results suggest that *scpr-1* is broadly required for generating the correct number of neuronal MTs, its function is conserved, and its overexpression can promote the formation of excessive MT polymers.

As an additional readout for the effect of SCPR-1 on MTs, we used a GFP::TBA-1/ α-tubulin transgene, which labels the MT lattice. Live imaging showed that the number of MT growth events was mildly reduced in *scpr-1* mutants compared to controls, consistent with the reduced number of polymers estimated by PTRN-1 labeling (**Figure S1H**). In the distal axon, where MT density is lower than in the proximal axon, loss of *scpr-1* led to the appearance of gaps between polymers, which could be rescued by expression of SCPR-1 in DA9 (**Figure 1F-H**). Estimation of average polymer length using an image analysis method, which we previously developed ^47^, showed a mild decrease in the dendrite (**Figure 1I, L**), and no change in the axon (**Figure S1K**), suggesting that the gaps are mostly due to reduced MT numbers rather than length. These results corroborate the observation made with PTRN-1::YFP, and suggest that the main defect in *scpr-1* mutants is a reduction in the number of MTs.

### *scpr-1* functions with *γ-tubulin* to promote microtubule nucleation

SCAPER was initially identified as an ER protein and later implicated in MT biology by several reports ^48–50^. Recently, it was shown to participate in tubulin autoregulation, a mechanism that destabilizes *tubulin* mRNA when soluble tubulin levels are experimentally raised by MT depolymerizing drugs ^51^. This prompted us to examine whether the phenotypes observed in *scpr-1* mutants are due to disrupted tubulin homeostasis. However, we were not able to detect differences in tubulin mRNA or protein levels in *scpr-1* mutants compared to control (**Figure S2A-C**), arguing that they do not underpin the reduction in MT polymer content in the mutant.

Since the specific reduction in polymer numbers in *scpr-1* mutants is suggestive of reduced MT nucleation, we next focused on γ-tubulin/TBG-1, the core component of the canonical MT nucleation complex γ-TuRC. We generated a cell-specific degradation allele for TBG-1 using CRISPR to tag its C-terminus with a ZF1 tag for degradation, and added a spGFP11 tag for visualization (referred to as *tbg-1 scKD –* see methods section for degradation details) (**Figure 2A**). Tagging did not affect TBG-1 function, as measured by viability and fertility of the animals when spGFP1-10 was ubiquitously expressed, and by reproducing the expected localization of TBG-1 in the germline (**Figure S3A-B**) ^52^. Cell-specific expression of the E3 ubiquitin ligase ZIF-1 in DA9 reduced TBG-1 levels below detection in ∼90% of animals, suggesting that *tbg-1(scKD)* represents a strong loss of γ-tubulin function (**Figure S3C-D**).

**Figure 2.**
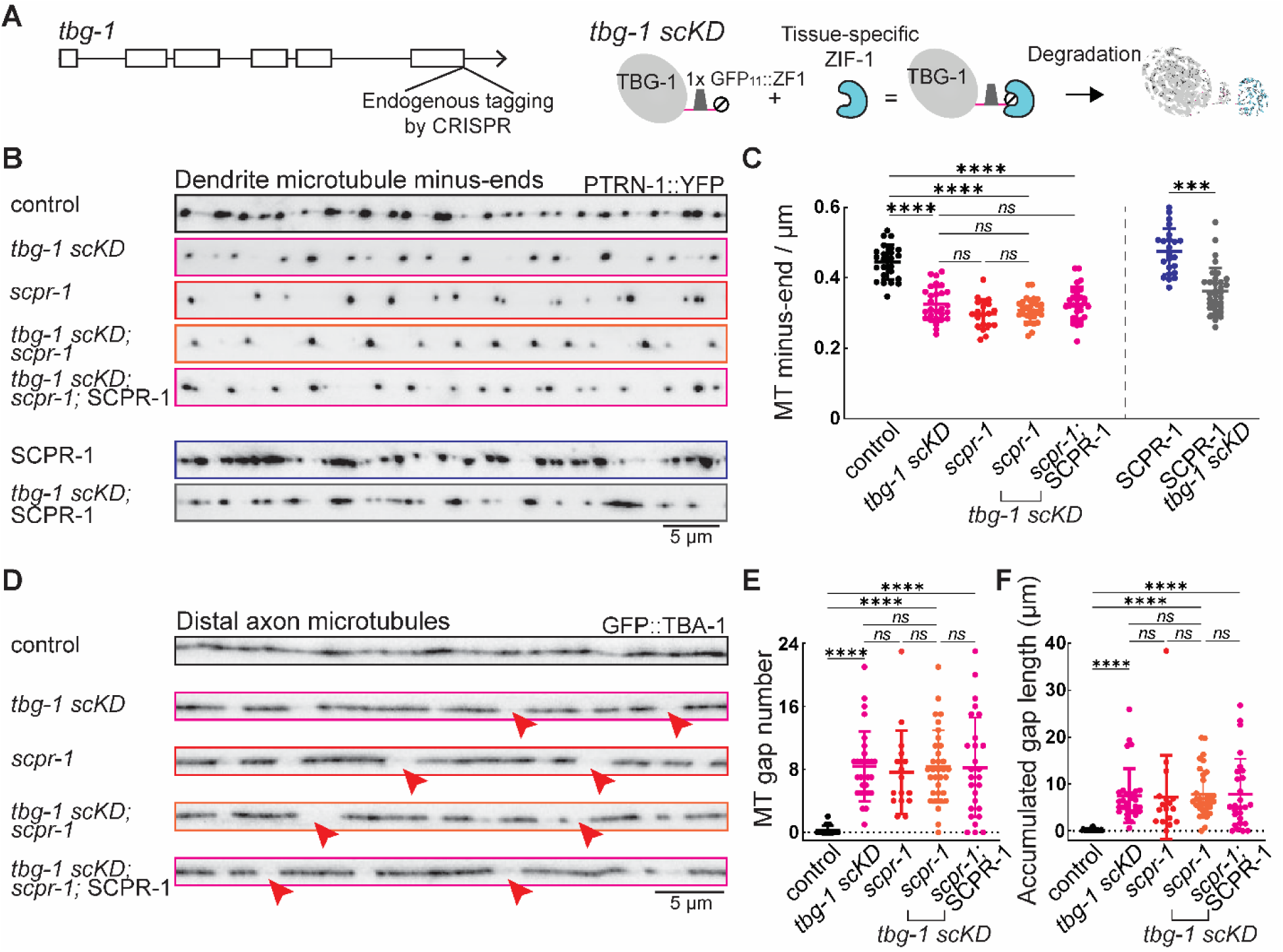
SCPR-1 functions with γ-tubulin to promote microtubule density. (A) Schematic of tbg-1 genomic region and the endogenously tagging technique. (B) Representative images of dendrite MT minus-end labeled by YFP-tagged PTRN-1 in control, *tbg-1 scKD*, *scpr-1*, *tbg-1 scKD; scpr-1* double mutant, SCPR-1 OE in *tbg-1 scKD; scpr-1* double mutant, SCPR-1 OE, and SCPR-1 OE in the *tbg-1 scKD* background, respectively. Scale bar = 5 μm. (C) Quantification of MT minus-end number density in DA9 dendrite in control (N=24), *tbg-1 scKD* (N=28), *scpr-1* (N=20), *tbg-1 scKD; scpr-1* double mutant (N=30), SCPR-1 OE in *tbg-1 scKD; scpr-1* double mutant (N=29), SCPR-1 OE (N=23), and SCPR-1 OE in the *tbg-1 scKD* background (N=33). ***: p<0.0005, ****: p<0.0001, and *ns*: not significant by Kruskal-Wallis tests. (D) Representative images of distal axon MT visualized by GFP-tagged MT subunit TBA-1 in control, *tbg-1 scKD*, *scpr-1*, *tbg-1 scKD; scpr-1* double mutant, and SCPR-1 OE in *tbg-1 scKD; scpr-1* double mutant, respectively. Red arrow heads indicate gaps between MT polymers due to MT mass reduction. Scale bar = 5 μm. (E and F) MT gap number and accumulated gap length in control (N=18), *tbg-1 scKD* (N=29), *scpr-1* (N=17), *tbg-1 scKD; scpr-1* double mutant (N=33), and SCPR-1 OE in *tbg-1 scKD; scpr-1* double mutant (N=27). ****: p<0.0001 and *ns*: not significant by Kruskal-Wallis tests.

Degradation of TBG-1 reduced PTRN-1::YFP to levels equivalent to those observed in *scpr-1* mutants (**Figure 2B, C**). Furthermore, in double mutants, the severity of the phenotype was not enhanced, suggesting that *scpr-1* and *tbg-1* function in the same genetic pathway (**Figure 2B, C**). Expression of SCPR-1 cDNA in *scpr-1; tbg-1(scKD)* double mutants did not restore the number of PTRN-1 puncta to control levels; conversely, TBG-1 degradation reduced the excessive number of PTRN-1 puncta caused by SCPR-1 overexpression (**Figure 2B, C**). These results are consistent with a scenario in which *tbg-1* functions downstream of *scpr-1* in MT nucleation.

Next, we examined whether the same genetic interactions can be observed using GFP::α-tubulin/TBA-1 as a marker for MTs. Indeed, single and double mutant analysis, as well as expression of SCPR-1 in *scpr-1; tbg-1* double mutants showed similar gaps in MTs at the distal axon (**Figure 2D-F**). As an additional readout for γ-tubulin and SCPR-1 function we estimated the ratio of soluble/polymerized tubulin with FRAP experiments, reasoning that reduced nucleation should increase soluble tubulin levels. As expected, *tbg-1 (scKD)* mutants showed a mild increase in the mobile fraction of GFP::TBA-1. A similar increase was observed in *scpr-1* mutants and in double mutants (**Figure S3E, F**). Taken together, these results indicate that *scpr-1* functions with *γ-tubulin* in nucleating neuronal MTs.

### SCPR-1 is required for γ-tubulin/TBG-1 recruitment to endosomes

To determine how SCPR-1 functions in γtubulin-dependent MT nucleation, we visualized its localization in DA9 neurons. Endogenous TBG-1, tagged with 2xspGFP11, was expressed at low levels in L4 larvae. Along the axon and the dendrite, TBG-1 was at the limit of detection, only appearing as a faint signal under long exposure (**Figure 3A, S4A**). At the tip of the dendrite – where MT nucleation occurs in early development ^24^ – and in the cell body, TBG-1 was readily detected in punctate structures. Endogenous SCPR-1, labelled with 7xspGFP11, was also faint along the axon and dendrite (**Figure S4B, B’**), and showed clear punctate distribution in the cell body (**Figure 3B**). We did not observe SCPR-1 enrichment at the tip of the dendrite and therefore used this location as a negative control when assaying the effect of *scpr-1* mutants on TBG-1 distribution. In the cell body, a transgene expressing RFP::SCPR-1 mimicked endogenous SCPR-1 distribution and strongly colocalized with TBG-1 in punctate structures (**Figure 3C, D**). These results indicate that SCPR-1 and TBG-1 colocalize with each other.

**Figure 3.**
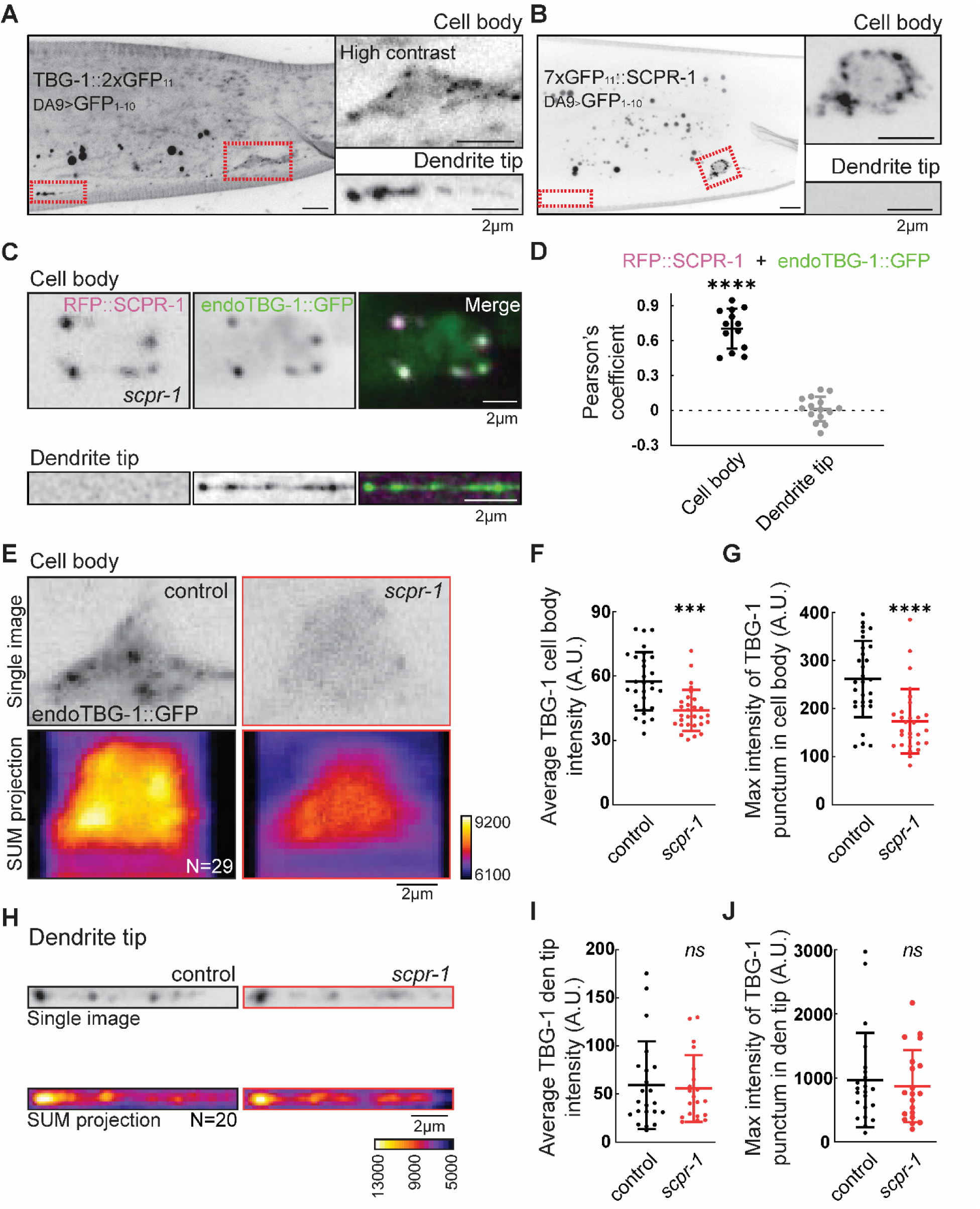
SCPR-1 is required for γ-tubulin recruitment to non-centrosomal MTOCs. (A and B) Representative images of endogenous labeled γ-tubulin/TBG-1 and SCPR-1, respectively, in C. elegans DA9 neuron. Red rectangles highlight cell body and dendrite tip regions. Enlarged images for cell body and dendrite tip regions were shown. Endogenous TBG-1 signal in cell body was increased contrast for visualization. Scale bar = 2 μm. (C) Representative images of RFP::SCPR-1 transgene expressed in the endogenous TBG-1::GFP with *scpr-1* mutant animal. Cell body and dendrite tip regions were shown. RFP::SCPR-1 was pseudo-colored as magenta, and endoTBG-1::GFP as green. Scale bar = 2 μm. (D) Plot of Pearson’s coefficient from RFP::SCPR-1 and endogenous TBG-1::GFP in cell body (N=13) and dendrite tip (N=15) regions. (E and H) Representative images of endogenous γ-tubulin signal in control and *scpr-1* mutant from cell body (E) and dendrite tip (H). Single images and SUM projection of multiple images pseudo-colored as heat map for cell body (N=29 for control and *scpr-1*) and dendrite tip (N=20 for control and *scpr-1*). Scale bar = 2 μm. (F and G) Plot of average intensity of γ-tubulin and maximum intensity of γ-tubulin punctum, respectively, in cell body from control (N=29) and *scpr-1* (N=29). ***: p<0.0005, and ****: p<0.0001 by Kruskal-Wallis tests. (I and J) Plot of average intensity of γ-tubulin and maximum intensity of γ-tubulin punctum, respectively, in dendrite tip from control (N=20) and *scpr-1* (N=20). *ns*: not significant by Kruskal-Wallis tests.

To determine the nature of the punctate structure containing SCPR-1, we examined its overlap with several compartment markers that were previously reported to colocalize with γ-tubulin. Endogenous SCPR-1 did not colocalize with RAB-11.1, RAB-5 or the Axin homolog PRY-1 (**Figure S5A-B**). In the cell body, SCPR-1 showed the strongest colocalization with the late endosomal marker RAB-7. Interestingly, in the axon, SCPR-1 colocalized poorly with RAB-7 and instead colocalized with RAB-6.2. In addition, expression of RAB-6.2 slightly influenced SCPR-1 distribution, causing it to appear more diffuse (**Figure S5C-H**). These results indicate that SCPR-1 localizes to vesicular endosomal compartments.

Next, we asked whether *scpr-1* is required for TBG-1’s punctate localization. We measured both the average TBG-1 intensity in the cell body, as well as the maximum intensity, which better reflects the accumulation in puncta. In *scpr-1* mutants, TBG-1 lost its punctate enrichment, appearing instead as a diffuse signal (**Figure 3E-G**). At the tip of the dendrite, where SCPR-1 is not detected, TBG-1 localization was unaffected in *scpr-1* mutants, arguing that the effect of SCPR-1 on TBG-1 is local (**Figure 3H-J**). *tbg-1* was not required for the localization of SCPR-1 (**Figure S4C, D**), consistent with the genetic interactions that suggest that SCPR-1 acts upstream of TBG-1. These results indicate that SCPR-1 acts at endosomal membranes to promote TBG-1 recruitment.

### SCAPER is required for γ-tubulin distribution and activity-dependent MT nucleation in mammalian neurons

We tested whether mammalian SCAPER performs a similar function to its *C. elegans* homolog, as suggested by the ability of human SCAPER to rescue the worm mutant (**Figure 1D, E**). Previous work established that in primary hippocampal neurons, γ-tubulin localizes to axonal *en passent* boutons and is required for activity-dependent MT nucleation at presynaptic sites ^34,53^ To test the role of SCAPER in this process, we delivered SCAPER shRNA into rat hippocampal neurons at DIV9, and measured MT growth and nucleation in these neurons at DIV16 by tagging growing MT plus-end with EB3-TdTomato. Consistent with previous findings, neuronal excitation evoked by application of the GABAA receptor antagonist bicuculline led to a robust increase in the number of EB3 comets (**Figure 4A-C**). SCAPER knockdown suppressed the effect of bicuculine on MT comet density (**Figure 4B-C**), suggesting that SCAPER is required for axonal MT nucleation upon neuronal activation.

**Figure 4.**
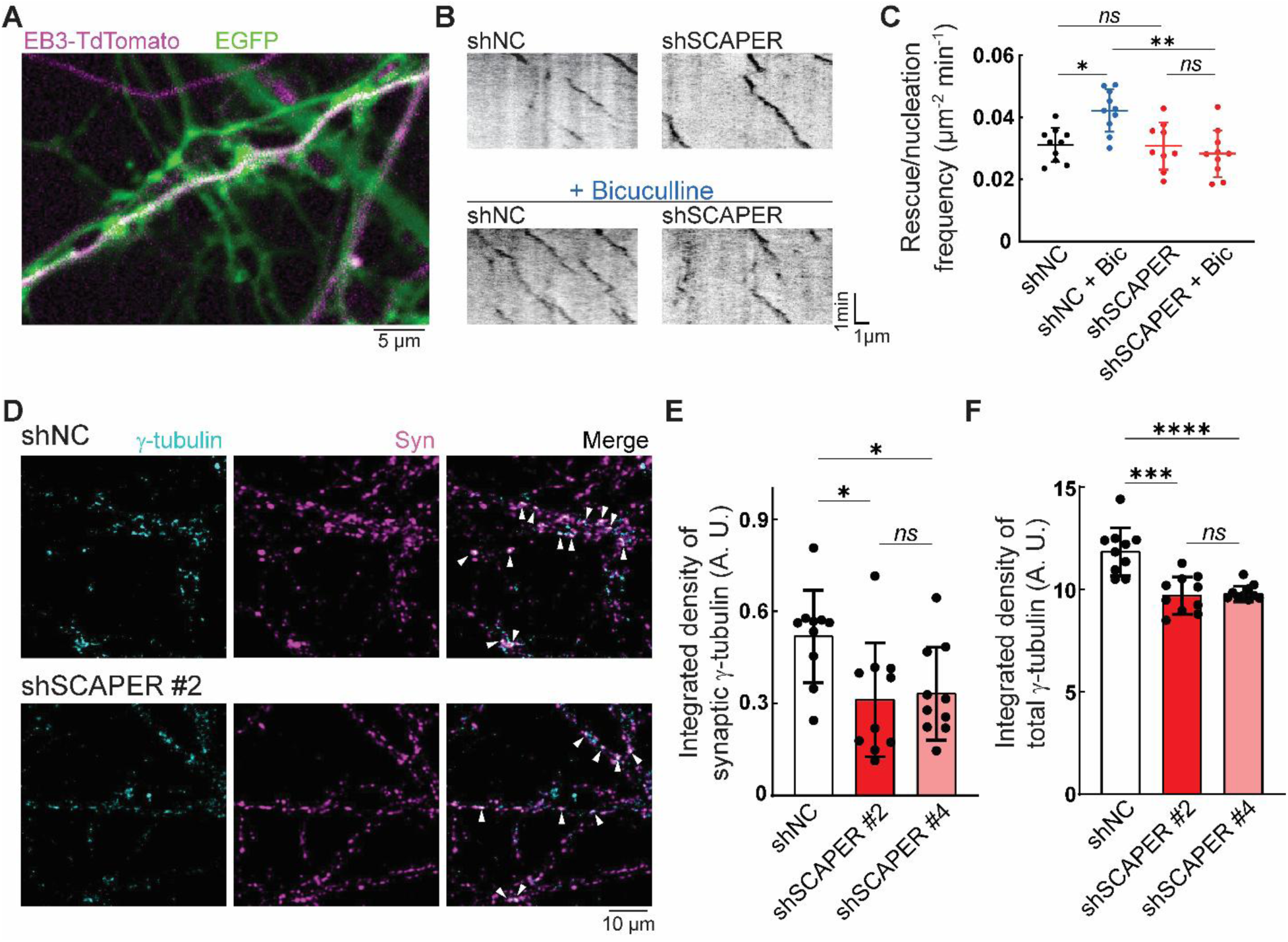
SCAPER is critical for γ-tubulin distribution and activity-dependent MT nucleation in mammalian neurons. (A) Representative images and relative kymographs of axons of hippocampal neurons expressing EB3-tdTomato after lentiviral delivery of shRNA non-coding control with EGFP reporter (shNC + Bic) or SCAPER with EGFP reporter (shSCAPER + Bic), for 7 days upon neuronal activation after incubation with bicuculline. Scale bar = 5 μm. (B) Representative kymograph of EB3 comets from shRNA negative control (shNC) and SCAPER shRNA knocking down in control and Bicuculline treatment. Scale bar = 1 min in time (y-axis) and 1 μm in length (x-axis). (C) Quantification of MT rescue/nucleation frequency in negative control of shRNA (shNC) and SCAPER shRNA (shSCAPER) in control and Bicuculline treated neurons. * (shNC vs shNC + Bic in C): p=0.0420. **: p<0.005, and *ns*: not significant by Kruskal-Wallis tests. (D) Maximum projection of spinning disk confocal images in WT hippocamapal neurons (14-16 DIV) fixed and immunostained for MAP2, γ-tubulin, and Synapsin 1/2. Neurons were infected with either a lentiviral non-coding control (shNC) or an shRNA targeting SCAPER (shSCAPER#2) with EGFP reporters for 7 days. White arrows indicate γ-tubulin signals co-localizing with synapsin 1/2. Scale bar = 10 μm. (E and F) Semiquantitative immunofluorescence of γ-tubulin intensity in hippocampal neurons treated as in (D). Total levels were measured as integrated density on the whole field of view, whereas synaptic γ-tubulin intensity was calculated by measuring the integrated density on the syn 1/2 mask (n=10 fields of view from N=2 independent experiments). * (shNC vs shSCAPER #2 in E): p=0.0115, * (shNC vs shSCAPER #4 in E): p=0.0185, ***: p<0.0005, ****: p<0.0001, and *ns*: not significant by Mann-Whitney tests.

We next examined whether SCAPER was also required for γ-tubulin distribution. Two independent SCAPER targeting shRNAs led to a reduction in γ-tubulin levels, an effect that was particularly evident at presynaptic sites (**Figure 4D-F, Figure S6A-B, E**), which in these neurons represent hotspots for γ-tubulin-mediated activity-dependent nucleation. Colocalization analysis between γ-tubulin and the presynaptic marker synapsin1/2 revealed the presence of γ-tubulin in a higher number of synapses in the knockdowns, while the total levels of synaptic γ-tubulin were reduced, suggestive of a failure to cluster γ-tubulin at specific synaptic sites (**Figure S6C-D**). These results indicate that mammalian SCAPER is required for promoting γ-tubulin-mediated activity-dependent nucleation in mammalian axons. This function is highly reminiscent of the role we identified for SCPR-1 in *C. elegans;* however, it is worth noting that mammalian SCAPER only promotes nucleation, as it is triggered by synaptic activity.

### SCPR-1 instructs γ-tubulin/TBG-1 recruitment for MT nucleation

The colocalization between TBG-1 and SCPR-1 at endosomes suggests that they act as nucleating centers. To test this possibility, we used live imaging to visualize MT growth events emanating from SCPR-1 puncta. Endogenously tagged end-binding protein 2 (EBP-2) was recruited to RFP::SCPR-1 puncta, and showed comets directly originating from them (**Figure 5A-B**), consistent with the idea that they act as MT nucleation sites. In agreement with this interpretation, we found that co-localization between SCPR-1 and EBP-2 at puncta was lost in *tbg-1 scKD* mutants (**Figure 5C, D**), and the number of EBP-2 comets in the cell body was reduced by 37.8% in both *tbg-1 scKD* and *scpr-1* mutants (**Figure 5B**). Together, these results suggest that SCPR-1 puncta are γ-tubulin-dependent MT nucleation sites.

**Figure 5.**
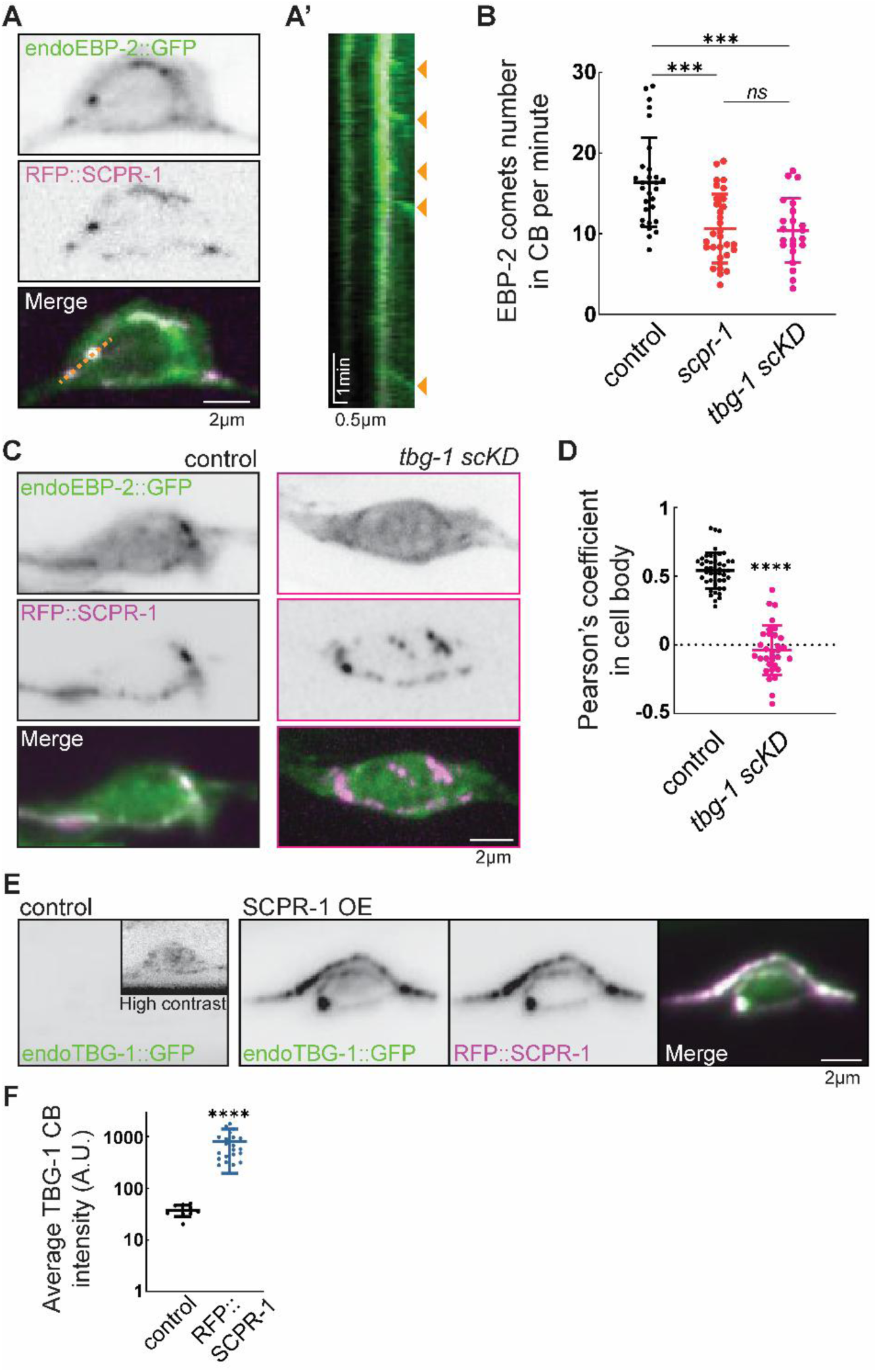
SCPR-1 directs γ-tubulin recruitment for MT nucleation. (A) Representative images of endogenous EBP-2 and RFP::SCPR-1 transgene. RFP::SCPR-1 and endoEBP-2::GFP were pseudo-colored as magenta and green, respectively. Yellow dashed line in merge channel was applied to generate kymograph in (A’). Scale bar = 2 μm. (A’) Kymograph of endogenous EBP-2 comets emerged from RFP::SCPR-1 puncta. Yellow arrow heads indicate EBP-2 comets from nucleation events. Scale bar = 1 min in time (y-axis) and 2 μm in length (x-axis). (B) Quantification of EBP-2 comets produced in DA9 cell body from control (N=30), *scpr-1* (N=31), and *tbg-1 scKD* (N=22). ***: p<0.0005 and *ns*: not significant by Kruskal-Wallis tests. (C) Representative images of endogenous EBP-2 and RFP::SCPR-1 transgene from control and *tbg-1 sckD*. RFP::SCPR-1 and endoEBP-2::GFP were pseudo-colored as magenta and green, respectively. Scale bar = 2 μm. (D) Plot of Pearson’s coefficient calculated in control (N=44) and *tbg-1 sckD* (N=32) from DA9 cell body. (E) Representative images of endogenous γ-tubulin recruited by RFP::SCPR-1 overexpression. Endogenous γ-tubulin images with same contrast level were shown in control and SCPR-1 OE. Insert in the control endoTBG-1::GFP was increased contrast for visualization. For SCPR-1 OE, endoTBG-1::GFP and RFP::SCPR-1 were pseudo-colored as green and magenta, respectively. Scale bar = 2 μm. (F) Measurement of average γ-tubulin signal in cell body from control (N=10) and RFP::SCPR-1 overexpression (N=21).

Next, we examined whether SCPR-1 was instructive for the recruitment of TBG-1. Overexpression of SCPR-1 led to a drastic increase in TBG-1 intensity in the cell body, with endogenous TBG-1 levels at puncta increasing by ∼20x (**Figure 5E, F**). Hence, SCPR-1 can instruct the recruitment of TBG-1, in agreement with the ability of SCPR-1 overexpression to increase MT numbers.

### The autoregulation factor TTC-5 functions with SCPR-1 and γ-tubulin in MT nucleation

In the tubulin autoregulation pathway, SCAPER is recruited to ribosomes containing *tubulin* mRNA by TTC-5, which recognizes the nascent peptide. In turn, SCAPER recruits the CCR4-NOT complex to destabilize *tubulin* mRNA ^51^. To test the role of the autoregulation pathway in MT nucleation we first generated a *ttc-5* deletion and asked how this affects neuronal MT abundance. Interestingly, *ttc-5* deletion reduced the number of PTRN-1 puncta in a manner that was indistinguishable from *scpr-1* or *tbg-1 scKD* mutants (**Figure 6A, B**). We generated a single cell degradation allele for TTC-5, and found that, similar to SCPR-1, TTC-5 functions cell autonomously (**Figure S7A, B**). In addition, double mutants *scpr-1; ttc-5* showed a reduction in PTRN-1::YFP that was identical to the single mutants (**Figure 6A, B**). We conclude that *scpr-1* and *ttc-5* function in the same genetic pathway to control MT abundance.

**Figure 6.**
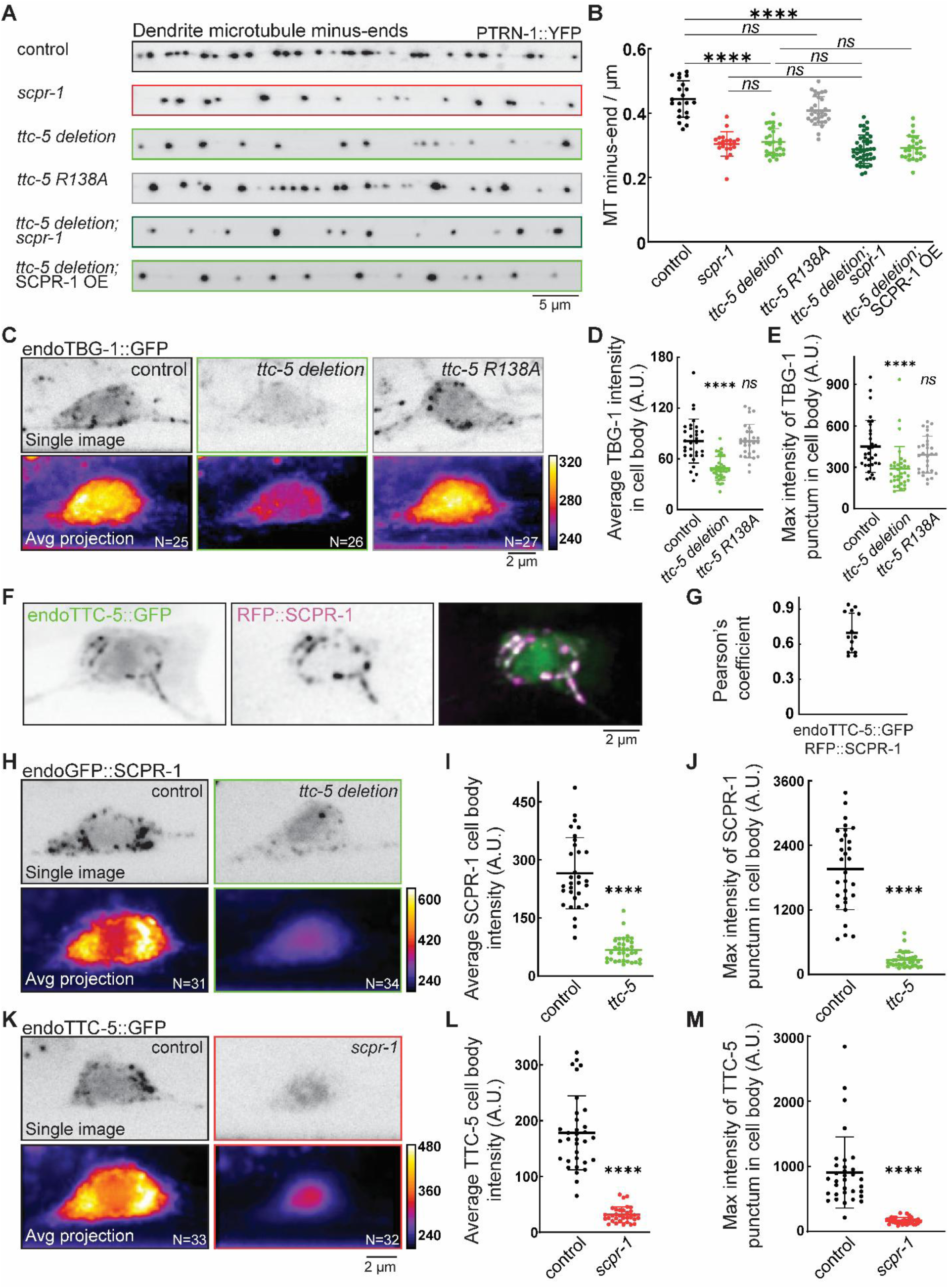
Tubulin autoregulation protein TTC-5 functions with SCPR-1 and γ-tubulin in MT nucleation. (A) Representative images of dendrite MT minus-end labeled by PTRN-1::YFP in control, *scpr-1*, *ttc-5 deletion; ttc-5 R138A*, and *ttc-5 deletion; scpr-1* double mutant. Scale bar = 5 μm. (B) Quantification of MT minus-end number density in DA9 dendrite in control (N=21), *scpr-1* (N=20), *ttc-5 deletion* (N=28), *ttc-5 R138A* (N=30), and *ttc-5 deletion; scpr-1* double mutant (N=42). ****: p<0.0001, and *ns*: not significant by Kruskal-Wallis tests. (C) Representative images of endogenous γ-tubulin signal in control and *ttc-5* mutants from cell body. Single images and average projection of multiple images pseudo-colored as heat map for cell body (N=25 for control, N=26 for *ttc-5 deletion*, and N=27 for *ttc-5 R138A*). Scale bar = 2 μm. (D and E) Plot of average intensity of γ-tubulin and maximum intensity of γ-tubulin punctum, respectively, in cell body from control (N=31), *ttc-5 deletion* (N=34), and *ttc-5 R138A* (N=29). ****: p<0.0001 and *ns*: not significant by Kruskal-Wallis tests. (F) Representative cell body image of RFP::SCPR-1 transgene expressed in the endogenous TTC-5::GFP animal. Endogenous TTC-5::GFP and RFP::SCPR-1 were pseudo-colored as green and magenta, respectively. Scale bar = 2 μm. (G) Plot of Pearson’s coefficient from RFP::SCPR-1 and endogenous TTC-5::GFP in cell body (N=14). (H) Representative images of endogenous SCPR-1 signal from control and *ttc-5 deletion* in cell body. Single images and average projection of multiple images pseudo-colored as heat maps for cell body were shown for control (N=31) and *ttc-5 deletion* (N=34). Spectrum bar shown as intensity [A. U.]. Scale bar = 2 μm. (I and J) Plot of average intensity of endogenous SCPR-1 and maximum intensity of SCPR-1 punctum, respectively, in cell body from control (N=31) and *scpr-1* (N=34). ****: p<0.0001 by Kruskal-Wallis tests. (K) Representative images of endogenous TTC-5 signal from control and *scpr-1* in cell body. Single images and average projection of multiple images pseudo-colored as heat maps for cell body were shown for control (N=33) and *scpr-1* (N=32). Spectrum bar shown as intensity [A. U.]. Scale bar = 2 μm. (L and M) Plot of average intensity of endogenous TTC-5 and maximum intensity of TTC-5 punctum, respectively, in cell body from control (N=33) and *scpr-1* (N=32). ****: p<0.0001 by Kruskal-Wallis tests.

Next, we tested the effect of TTC-5 on SCPR-1 and TBG-1. Loss to TTC-5 led to a strong reduction in γ-tubulin recruitment to endosomal puncta (**Figure 6C-E**). In addition, TTC-5 colocalizes with SCPR-1 at the endosomal puncta, suggesting that they act together locally (**Figure 6F, G**). In agreement, SCPR-1 and TTC-5 were required for each other’s enrichment at the puncta (**Figure 6H-M**). These results indicate that two proteins from the tubulin autoregulation pathway, SCPR-1 and TTC-5, function together with γ-tubulin to promote MT nucleation.

In the autoregulation pathway, SCPR-1 recruits the CCR4-NOT complex to destabilize tubulin mRNA ^51^. We tested the effects of *ttc-5* deletion on tubulin mRNA, and found no effect, suggesting that TTC-5 promotes MT nucleation independently of autoregulation (**Fig S2A**). These results suggest that upstream components of the autoregulation pathway, TTC-5 and SCPR-1, participate in γ-tubulin-dependent MT nucleation, but downstream components that destabilize mRNA do not regulate nucleation. Consistent with this idea, CRISPR-generated scKD alleles for the CCR4-NOT subunits *ccr-4* and *not-11* did not lead to a reduction in PTRN-1 puncta and did not colocalize with SCPR-1 (**Fig S6A, C-K**).

Last, to genetically distinguish the roles of TTC-5 in autoregulation from nucleation, we generated a point mutation in a conserved arginine that is essential for its interaction with the nascent tubulin peptide. An R147A mutation (corresponding to R138A in *C. elegans*) in TTC5 completely blocks autoregulation ^54^. In contrast to the *ttc-5* deletion, the *ttc-5 (R138A)* allele did not reduce PTRN-1 puncta (**Figure 6A-B**) and had no effect of TBG-1 recruitment to endosomes (**Figure 6C-E**). Thus, TTC-5 function in nucleation is independent of its role in autoregulation.

### SCPR-1 promotes axon regeneration by recruiting TBG-1 to the growth cone

MTs are essential for axon regeneration. In *C. elegans,* MT dynamics increase 2-3 hours after axotomy ^55–57^, raising the possibility that MT nucleation may play a critical role during this time window. To test the role of γ-tubulin and SCPR-1 in this process, we performed laser axotomy of the DA9 axon at the level of the commissure (**Figure 7A**). We quantified the percentage of axons which successfully regenerated and the extent of axon regrowth, as previously described ^58^. 24 hours after axotomy, ∼58% of control axons regenerated successfully. In comparison, in *scpr-1* mutant regeneration was strongly impaired, with only ∼20% of axons regenerating successfully (**Figure 7B, C**). *scpr-1* was also required for the regeneration of GABA motor neurons (**Figure S8A-C**), indicating that its function is not restricted to DA9. In *tbg-1 scKD* mutants and *scpr-1; tbg-1 scKD* double mutants, regeneration rates were reduced to the same extent as in *scpr-1* single mutants, indicating that *scpr-1* and *tbg-1* function in the same genetic pathway to promote axon regeneration (**Figure 7B-D**). Measurements of axon regrowth length showed that the single and double mutants were significantly impaired in growth compared to control axons (**Figure 7D**), suggesting that SCPR-1 and TBG-1 function in axon regeneration by promoting regrowth of injured axons.

**Figure 7.**
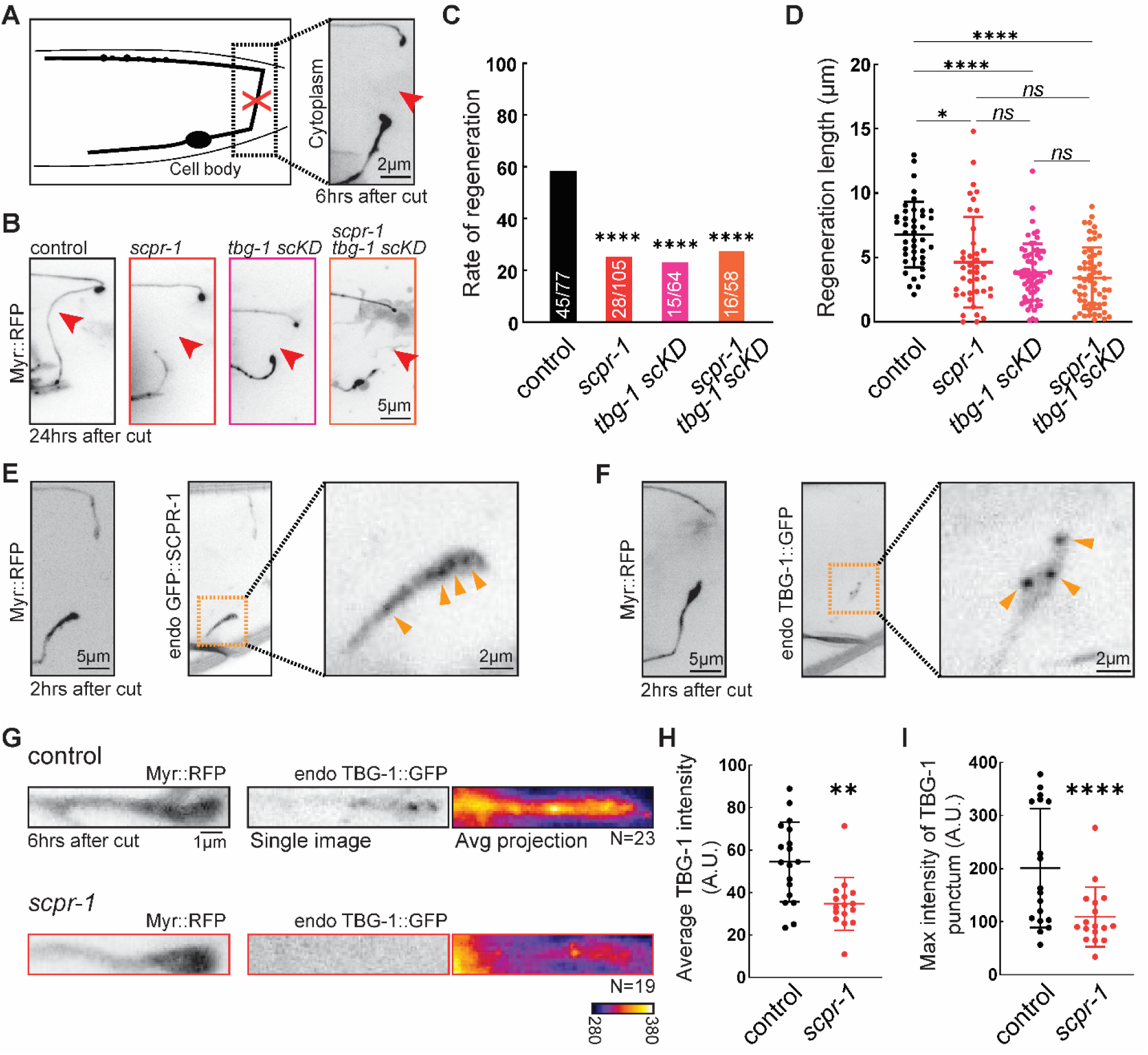
SCPR-1 recruits γ-tubulin to growth cone to promote axon regeneration. (A) Schematic of axotomy in DA9. One representative image of plasma membrane marker of DA9 was shown 6 hours after axotomy. Red cross indicates the position of cutting. Red arrowhead highlights regrowth of axon after cutting. Scale bar = 2 μm. (B) Representative images of successful regeneration from control, and unsuccessful regrowth from *scpr-1*, *tbg-1 scKD*, and *scpr-1; tbg-1 scKD* double mutant, 24 hours after cut. Scale bar = 5 μm. (C) Quantification of regeneration rate in control (N=45/77), *scpr-1* (N=28/105), *tbg-1 scKD* (N=15/64), and *scpr-1; tbg-1 scKD* double mutant (N=16/58). Two-sided Fisher’s exact tests were applied. ****: p<0.0001. (D) Quantification of regrowth length from control (N=41), *scpr-1* (N=41), *tbg-1 scKD* (N=63), and *scpr-1; tbg-1 scKD* double mutant (N=58), 24 hours post axotomy. Brown-Forsythe and Welch ANOVA tests were applied. * (*scpr-1* vs control): p=0.0134, ****: p<0.0001. *ns*: not significant. (E and F) Representative images of endogenous SCPR-1 (E) and TBG-1 (F) recruited to growth cone 2 hours after axotomy. Plasma membrane and endogenous GFP images were shown. Enlarged growth cone regions highlighted by red rectangles were shown. Scale bar = 5 μm. (G) Representative images of endogenous γ-tubulin signal from control and *scpr-1* in growth cone along with plasma membrane marker 6 hours post axotomy. Single images and average projection of multiple images pseudo-colored as heat maps were shown for control (N=19) and *scpr-1* (N=23). Spectrum bar shown as intensity [A. U.]. Scale bar = 1 μm. (H and I) Plot of average intensity of endogenous TBG-1 (H) and maximum intensity of TBG-1 punctum (I), respectively, from growth cone in control (N=18) and *scpr-1* (N=16). Kruskal-Wallis tests were applied. **: p<0.005 and ****: p<0.0001.

To gain insight into the function of SCPR-1 and TBG-1 in promoting axon regrowth, we imaged growth cones 2 hours post axotomy, when MT dynamics are elevated during regeneration. We found that both SCPR-1 and TBG-1 were clearly present at regenerating growth cones, at levels higher than their normal axonal localization, suggesting that they function locally (**Figure 7E, F**). Reminiscent of its function in the cell body, we found that *scpr-1* was required for the enrichment of TBG-1 on punctate structures in the growth cone during axon regrowth (**Figure 7G-I**). These results suggest that SCPR-1 functions in axon regeneration by enriching γ-tubulin at the growth cone to promote axon regrowth.

Taken together, our results identify a role for the tubulin autoregulation factors TTC-5 and SCPR-1 in γ-tubulin mediated MT nucleation, where they function by recruiting TBG-1 to endosomes to control neuronal MT content during homeostasis and to promote robust axon regrowth during regeneration.

## Discussion

How neurons establish a cytoskeletal architecture composed of the correct number of MT polymers is unclear. Here we found that two proteins from a pathway that controls *tubulin* mRNA stability promote non-centrosomal MT nucleation in neurons through γ-tubulin. *scpr-1, ttc-5* and *tbg-1/γ-tubulin* function in the same genetic pathway, which accounts for ∼30% of the MT content in the *C. elegans* motor neuron DA9. The three proteins colocalize with each other, and both SCPR-1 and TTC-5 promote the recruitment of γ-tubulin to endosomal compartments where nucleation occurs. SCPR-1 can also instruct excessive γ-tubulin recruitment, and its overexpression increases neuronal MT content. Mammalian SCAPER likely has related functions, because it can rescue *C. elegans scpr-1* mutants, and rat SCAPER is required for activity-dependent MT nucleation in axons and the accumulation of γ-tubulin at presynaptic sites. Finally, *C. elegans* SCPR-1 and TBG-1/γ-tubulin are not required for normal axon elongation, but function in the growth cone of regenerating axons to promote axon regrowth after injury. This study uncovers regulatory mechanisms that underpin the formation and function of the neuronal MT cytoskeleton and reveal an unexpected overlap between two pathways that regulate cytoskeletal homeostasis: autoregulation and nucleation.

γ-Tubulin Ring Complex nucleates MTs in most, but not all, cells studied to date ^59–61^. Since different cell types vary considerably in their MT architecture, mechanisms of γ-tubulin recruitment, orientation, and activation are cell-type specific. In neurons, γ-tubulin and its associated proteins have been shown to function at centrosomes in early development and at non-centrosomal sites at later stages^27,30,62^. Recent work identified neuronal non-centrosomal MTOCs at Golgi, Golgi outposts. Ciliary base and various endosomal populations ^22,24–26,63,64^; however, how these MTOCs form remains ill defined. We found that recruitment of γ-tubulin to non-centrosomal MTOCs requires SCPR-1 and TTC-5 and is necessary for MT nucleation at these sites and the formation of the neuronal MT array. Interestingly, in the axon, SCPR-1 colocalized better with RAB-6.2 than with RAB-7. RAB6 regulates both retrograde and anterograde trafficking. In the cell body, it localizes to the Golgi, while in neurites, it is on motile vesicular structures that transport cargo such as endocytosed AMPA receptor GLR-1 ^65–67^. Recent work suggests that RAB6 vesicles are captured at presynapses by ELKS, which bears homology to Golgi proteins that mediate vesicle tethering at the Golgi ^68^. This is possibly related to the fact that both Golgi outposts and presynaptic markers have been identified as sites of non-centrosomal MT nucleation and may also explain why rat SCAPER specifically promotes the γ-tubulin enrichment and activity-dependent nucleation at *en passan*t boutons. However, testing these scenarios would require further studies, since SCAPER levels were too low to detect with immunofluorescence in rat neurons, and in *C. elegans,* the expression of RAB-6.2 had a mild effect on SCPR-1 distribution.

The relative contributions of different nucleation mechanisms to the overall MT density and to neuronal morphogenesis are unclear. Since γ-tubulin is essential for cell division, it is challenging to completely eliminate it to assess its roles in early axon development in vivo. We degraded γ-tubulin to undetectable levels by the time the axon is still elongating rapidly with the growing animal’s body and found no effect on axon length. The reduction in MT density in *scpr-1, ttc-5,* and *tbg-1 scKD* mutants was ∼30%, which, together with other studies, suggests the existence of redundant nucleation mechanisms ^69,70^. It is also possible that in the axon, a high tubulin concentration reduces the barrier to nucleation, allowing various MAPs to compensate for the loss of γ-tubulin. In comparison, we observed a strong phenotype when we examined axon regrowth after regeneration in *scpr-1* or *tbg-1 scKD* mutants. This suggests that γ-tubulin-mediated nucleation is essential when a burst of localized nucleation is required to support growth.

Our results reveal an intriguing connection between MT nucleation and autoregulation. In *C. elegans,* several observations indicate that autoregulation does not determine neuronal MT content: First, if SCPR-1 and TTC-5 functioned in autoregulation, one would expect their loss to increase, rather than decrease, MT abundance. Second, we found no difference in tubulin mRNA or protein levels in the mutants. Third, *ccr-4* and *ntl-11* were not required for neuronal MT density, and thus, only upstream factors from the autoregulation pathway are involved. The involvement of SCAPER in MT nucleation is likely not limited to *C. elegans* since human SCAPER rescued the *C. elegans* mutant. Furthermore, we found that mammalian SCAPER is required for synaptic microtubule nucleation and γ-tubulin distribution, highly reminiscent of SCPR-1 function. Since we were unable to visualize rat SCAPER with two commercially available antibodies, we could not determine whether it colocalizes with γ-tubulin, and thus whether it acts directly, as it does in *C. elegans*, remains to be determined. Together, our data raises the possibility that autoregulation and nucleation might be related or serve similar cellular purposes. In particular, since the signals that trigger autoregulation are unknown, it is tempting to speculate that nucleation might be involved. However, establishing such a direct link between these two pathways would require further work.

Last, both SCAPER and TTC5 have been initially identified in contexts unrelated to either autoregulation or nucleation. SCAPER was initially identified as an ER protein that is regulated in a cell-cycle-dependent manner ^48^, and TTC5 was identified as a nuclear protein interacting with the acetyltransferase p300 ^71^. Subsequent work identified a role for TTC5 in inhibiting actin nucleation ^72^, while several reports implicate SCAPER in MT biology, regulating cell division and cilia length ^73–75^. Recently, TTC5 and SCAPER were shown to interact with each other and promote the recruitment of the CCR4-Not complex to *tubulin-*translating ribosomes ^38^. Our results support a role for TTC5 and SCAPER in MT nucleation, a function that is consistent with the phenotypes of patients with mutations in the proteins. In humans, mutations in SCAPER, TTC5 and γ-tubulin/TUBG1 are associated with a phenotypic spectrum that is highly suggestive of a role in neurodevelopment ^76–78^. It is interesting to note that some phenotypes caused by TUBG1 mutations, such as posterior predominant pachygyria/lissencephaly, are suggestive of a role in neuronal migration, but are absent in patients with TTC5 and SCAPER mutations, which may be related to their specific role in non-centrosomal MT nucleation, whereas TUBG1 functions both in centrosomal and non-centrosomal nucleation. In summary, our work reveals an unexpected role for tubulin autoregulation factors in neuronal microtubule nucleation, providing insight into how neurons assemble their cytoskeletal architecture.

## Acknowledgements

The authors would like to thank the Yogev lab for fruitful discussions and feedback on this work. Some strains were provided by the CGC, which is funded by NIH office of Research Infrastructure Programs (P40 OD010440). This project is funded by NIH grant R35GM133573 to S.Y, R01NS094219 to M.H and R01AG050658 to F.B.

## Materials and Methods

### Stains and maintenance

*C. elegans* strains were cultured on nematode growth medium (NGM) plates seeded with OP50 bacteria following standard procedures. Cultivation temperature was 20°C unless otherwise indicated. Stains used in this study are listed in Table S1.

### Experimental models

All protocols and procedures for rats were approved by the Committee on the Ethics of Animal Experiments at Columbia University and were conducted in accordance with the Guide for the Care and Use of Laboratory Animals, published by the National Institutes of Health. Time-pregnant Sprague Dawley rats (Embryonic Day 18) were purchased from Charles River Laboratories for primary hippocampal neuronal cultures. All protocols and procedures for mice followed the National Institutes of Health guidelines and were approved by the Institutional Animal Care and Use Committee of Columbia University.

### Genetic screen

*scpr-1(shy5)* allele was isolated from a forward EMS mutagenesis screen for mutants with reduced PTRN-1::YFP puncta in DA9. Mutagenesis was performed as previously described and screening was done on compound fluorescence microscope. Mutants were outcrossed x4 and mapped using the “Sibling subtraction method” ^79^. Standard procedures were used for DNA purification, Illumina sequencing and sequence alignments.

### Gene-editing by CRIPSR/Cas-9

All CRISPR-based gene-editing experiments were performed according to the protocols from Mello lab and a recently updated single-strand DNA as the repair template ^80,81^. In brief, 0.5 μL Cas9 (S. pyogenes Cas9 Nuclease V3, 10 μg/ μL, IDT catalog # 1081058), 2-3 μg gRNA (1 μg for each if two gRNAs were used), 20-40 pmol dsDNA or 2-6 pmol ssDNA repair template with 30bp homology arms, 800 ng pRF4[rol-6(su1006)] plasmid as a selection marker, and nuclease free water were added to reach total volume of 20 μL. The NGG PAM sites were selected using Benchling (benchling.com) with 20 bp length and selected by > 55 on-target score and > 50% GC content. The gRNAs were synthesized from DNA oligos using the EnGen sgRNA synthesis kit (NEB catalog #E3322S) and subsequently purified by Monarch RNA cleanup kit (NEB catalog # T2040). The dsDNA repair template was melted and cooled before adding to the injection mix. In case of preparing ssDNA repair template, 5’ phosphate was added to one of the oligo strands which was treated by Lambda exonuclease (NEB catalog #M0262S) for degradation. The purified ssDNA was added to the injection mix. The mixture was centrifuged at 13,000 rpm for 20 min and the supernatant was kept on ice and used for injection. F1 rollers were singled and genotyped to screen for the correct editing events. The homozygotes were verified by Sanger sequencing and outcrossed at least 2 times. All the gene-editing strains generated in this study are listed in Table S2.

### Cloning and transgenes

Plasmids were generated by Gibson Assembly and verified by Sanger or Nanopore sequencing. At least 3 transgenic lines were selected from each injection. For *mig-13* promoter driven expression, 3-10 ng/μL were used, and 24-28 ng/μL for plasmids containing *mig-13* with a LoxP-flanked STOP cassette were injected to *unc-4c* promoter driven CRE expression strain. A list of all plasmids used or generated in this study is shown in Table S3. All sequences and plasmids are available upon request from the corresponding author.

Extrachromosomal arrays were integrated into the genome by 300 μJoules 365nm UV treatment after 15 min incubation with 1mL of 30 μg/μL Trimethylpsoralen (TMP, Thermo Fisher catalog # J63226.03). F1s with stable expression of co-injection markers were selected, mapped, and outcrossed at least 5 times.

### Cell specific expression and degradation

We expressed transgenes in DA9 using *mig-13* promoter, which also turnes on in neurons overlapping with DA9, and pharyngeal cells required for viability. To limit narrow the expression pattern of *mig-13* promoter, we used a binary system with an *unc-4c* promoter driven CRE expression in DA9 but not VA12 or pharyngeal cells, and a LoxP-flanked stop cassette before ORF ^82^.

Single cell degradation of protein-of-interest was achieved by the ZF1-ZIF-1 system or auxin-inducible TIR1 ^83,84^. ZF1 or AID degron was endogenously tagged to the gene-of-interest. To avoid lethality from degrading γ-tubulin/TBG-1 in pharyngeal cells, we used the CRE-LoxP system described above to prevent expression of ZIF-1 in these cells. For auxin-induced degradation, TIR1 was expressed under *mig-13* promoter. L1 animals were transferred from regular NGM plates to plates containing 1 mM Auxin (Alfa Aesar catalog #A10556) and one day-old-adults (1 doa) were mounted for imaging.

### Fluorescence microscopy

For still imaging, L4 or 1 day old adults animals were incubated in 10 mM levamisole diluted in M9 on a 2% agarose pad. For time-lapse live-imaging, L4 worms were incubated in 0.5 mM levamisole diluted in M9 for 20 min or until paralysis and transferred onto a 10% agarose pad with a droplet of M9. All images were acquired with DMi8 inverted microscope (Leica) equipped with a VT-iSIM system (BioVision) and an ORCA-Flash 4.0 camera (Hamamatsu).

Still and time-lapse images were acquired by MetaMorph under HC PL APO 63x/1.40NA OIL CS2 or HC PL APO 100×/1.47NA OIL objectives. Images were taken with 25% laser power with 200 or 300 ms exposure time, except for endogenously labeled proteins which were imaged with 2-3 seconds exposure. FRAP experiments were performed by Visitron system with 405 nm laser using VisionView. A box of ∼20×80 px region were chosen to bleach, and time-lapse images were documented every 1s. 3 images before FRAP were recorded for normalization.

### Image editing and quantification

Raw microscopy images were edited by FIJI/ImageJ. Still images with z stacks were processed by maximum intensity projection. For images covering the whole DA9 neuron, multiple images were assembled by the pairwise-stitching plugin with linear blending fusion.

The PTRN-1::YFP number was measured by the BAR Find Peaks plugin and divided by the length of the dendrite/axon to calculate the density of microtubule minus-ends. The gaps between GFP::TBA-1 signals were measured using line segments. A matlab algorithm MT-Quant was applied to simulate the microtubule mass (Cooper *et al.*, 2016; Yogev *et al.*, 2016).

Kymographs were made by the KymoResliceWide plugin in ImageJ for time-lapse images. Microtubule dynamic events were defined as displacement of GFP::TBA-1 signal of greater than 0.5 μm, where one side shits but the other remains stationary.

For FRAP data, one non-FRAP region was measured and used for normalization to minimize potential effects of focal plane or biological drift. Normalized intensity = (FRAPed region intensity - background) / (non-FRAPed region intensity - background). The mobile fractions = (Y plateau – Y0) / (Y-1 - Y0). Y-1 is the average intensity of the ROI from 3 timepoints before FRAP.

For colocalization analysis, cell body regions were selected for Pearson coefficient calculation by ImageJ Coloc two plug-in using Costes threshold regression with PSF = 1.0 and Costes randomization = 10.

All quantifications were conducted on raw images.

### Primary hippocampal neuronal cultures

Primary hippocampal neuronal cultures were prepared as previously described ^85^. Briefly, hippocampi were dissected from E18 rats and pooled together, and neurons were plated at a density of 7 × 10^4^ cells/dish on 100 μg/mL poly-D-lysine-coated 12-well plates for live imaging, immunostaining, mRNA extraction, and quantification in 35 mm MatTek live imaging dishes. Primary neurons were maintained in Neurobasal medium (Invitrogen) supplemented with 2% B-27 (Invitrogen) and 0.5 mM glutamine (Invitrogen) at 37°C with 5% CO_2_.

### Lentiviral shRNA silencing

Production of lentiviral particles was conducted using the 2^nd^ generation packaging system as previously described ^85^. Briefly, HEK293T cells were co-transfected with lentiviral shRNA constructs and the packaging vectors pLP1, pLP2, and pLP-VSV-G (Invitrogen) using calcium phosphate. 48 h and 72 h after transfection, the virus was collected, filtered through 0.45 µm filter, and 1 volume of cold Lentivirus Precipitation Solution (Alstem) was added to every 4 volumes of lentivirus-containing supernatant and incubated for at least 6 h at 4°C. The lentivirus-containing supernatant was centrifuged for 30 minutes at 3000 rpm, and the pellet was resuspended in the minimum volume of neuronal medium recommended by the manufacturer. The concentrated virus was aliquoted and stored at −80°C. Lentiviral constructs to knockdown rat SCAPER in hippocampal neurons were purchased from VectorBuilder with the following target sequence shSCAPER#2: GAATGATGTTCTCGCAGATTA and shSCAPER#4: ATACACCAATTGACGGTTTAT. The vector with the mammalian noncoding (shNC) sequence was used as a control. Both SCAPER and shNC shRNA vectors were driven by the human U6 promoter, with an EGFP reporter gene expressed under the hPGK promoter in all constructs.

### AAV expression of tagged MT plus-end marker

AAV virus to express EB3-tdTomato was purchased from Vectorbuilder.

### Live imaging of MT dynamics in hippocampal neuronal cultures

Lentiviral delivery of SCAPER shRNA was performed in hippocampal neurons at 7 DIV. The next day, neurons were co-infected with AAV expressing EB3-tdTomato at 8 DIV. For hippocampal neurons, live-cell imaging was performed at 14-16 DIV, 6-8 days after infection, in complete HBSS media using an IX83 Andor Revolution XD Spinning Disk Confocal System. The microscope was equipped with a 60 x /1.35 oil UPlanSApo objective, a multi-axis stage controller (ASI MS-2000), and a controlled temperature and CO2 incubator. Movies were acquired with an Andor iXon Ultra EMCCD camera and Andor iQ 3.6.2 live cell imaging software at 2 s/frame for 3 min. To induce neuronal activity, neurons were washed 1x with complete HBSS media and 20 µM bicuculline or DMSO control was added to complete HBSS media after washes. Movies were taken 1 minute after treatment. Axons were selected during live imaging based on morphology and the exclusive anterograde movement of EB3 comets. Confocal live imaging acquisition was performed by acquiring three stacks of 0.7 µm each, and the maximum projection of the movies was analyzed using ImageJ. Axons were selected during live imaging based on morphology and the exclusive anterograde movement of EB3 comets. Kymographs were generated by drawing a region in the distal axons (more than 100 µm from the cell body) based on morphology and anterograde movement of EB3-labeled comets. Parameters describing MT dynamics were defined as follows: rescue/nucleation frequency: number of rescue or nucleation events per μm^2^ per min; catastrophe frequency: number of full tracks/total duration of growth; comet density: number of comets per μm^2^ per min; growth length: comet movement length in μm; comet lifetime: duration of growth; growth rate: growth length/comet lifetime ^86,87^.

### Immunofluorescence microscopy and analyses

Neurons were fixed in 4% PFA + 4% sucrose for 20 min at RT. Cells were then washed in PBS, permeabilized with 0.2% Triton X-100 for 10 min, blocked in 3% BSA in PBS for 1 h, and stained with primary antibodies for 2 h followed by secondary antibodies for 1 h. Donkey anti-chicken AlexaFluor 405 (A48260), donkey anti-rabbit AlexaFluor 594 (A21207), and donkey anti-mouse AlexaFluor 647 (A31571) (Invitrogen) secondary antibodies were used for immunostaining of MAP2, synapsin 1/2 and γ-tubulin, respectively. Samples were kept in live cell dishes and mounted with Ibidi mounting medium. Samples were observed with the same spinning disk confocal setup that was used for live imaging at 60x magnification.

All images were analyzed by ImageJ software. Analysis of colocalization was performed by measuring colocalization between γ-tubulin and synapsin 1/2 using JaCOP colocalization plugin. Semiquantitative IF analysis of γ-tubulin intensity was performed by plotting the integrated density of the total γ-tubulin signal or only the synaptic γ-tubulin signal, obtained by measuring the integrated density of the γ-tubulin signal after creating a mask on the synapsin 1/2 signal.

### SCAPER mRNA expression analysis

Total RNA was isolated from hippocampal neurons using TRIzol reagent (Invitrogen). Synthesis of first-strand cDNA was performed by reverse transcription of total RNA using SensiFAST cDNA Synthesis Kit (Bioline, London, UK) according to the manufacturer’s protocol. The Mx3000P Real-Time PCR System (Agilent-Stratagene) was employed to perform quantitative real-time PCR analysis (qRT-PCR) of the indicated mRNA expression levels. The reaction mix containing the cDNA template, the SensiFAST SYBR® Lo-ROX Kit (Bioline, London, UK) and the primer was amplified using standard qPCR thermal cycler parameters. Each sample was amplified in triplicate, and the quantification of the mRNA was performed using MxPro qPCR software. The average of the three threshold cycles was used to calculate the number of transcripts. Data were normalized with the endogenous housekeeping gene (HPRT) and expressed as the fold change with respect to the control sample value.

The following qRT-PCR assays were used:

SCAPER (for shSCAPER #2: rSCAPER forward, GGTATGTCTCTGGACTGGAATG; rSCAPER reverse, GCAGGCTCCTCTTCTACAATATC; for shSCAPER #4: rSCAPER forward, CCCAGCCACAAACATGAAC; rSCAPER reverse, GCAATCAGACAGCTAAAAACC); HPRT (rHPRT forward, CGCTGAATAGAAATAGTGATAGGTCC; rHPRT reverse, GACGTTCTTTCCAGTTAAAGTTGAG);

### Quantitative real-time PCR for *C. elegans* transcripts

The synchronized 1-day-old adult animals were washed with M9 and collected by centrifuging at 3000rpm for 1-2 min, followed by TriZol treatment (TRIzol LS Reagent, Thermo Fisher, catalog # 10296010) and RNA extraction. The RNA extracts were reverse transcribed (QuantiTech Reverse Transcription Kit, Invitrogen, catalog # 205311) and used for real-time PCR (QuantiTech SYBR Green RT-PCR, Invitrogen catalog # 204143; Applied Biosystems 7500 Fast Real-Time PCR System). Primers were designed by NCBI primer blast with a 115-125 bp target region at the C-terminus of the gene-of-interest and Tm temperature at 59-61°C. Relative expression levels were calculated based on the level of housekeeping gene *rpl-15*. Each transcript was normalized to the wild-type level or untreated animals with the same genetic background. Animals from at least 3 biological replicates were collected and applied for RT-PCR. The primers used for RT-PCR are listed in Table S4.

### Laser axotomy

The axotomy was performed with Micropoint pulsed UV laser system attached to a Nikon Eclipse 80i microscope, with detailed parameters described previously ^88^. L4 stage worms were mounted with 0.05-micron Polybead Microspheres (catalog #08691-10) between 3% agarose pad on a slide and a No. 1.5 coverslip. DA9 and GABA neuron axons were identified by cell shape markers under 100x Plan Apo VC lens (1.4 NA) and severed at the midpoint of commissures. Injured animals were then allowed to recover on OP50 plates for later imaging.

### Immunoblotting

1 day-old *C. elegans* were washed and collected by M9. The pallet of worms froze immediately by using liquid nitrogen. Total amount of proteins was tested by the Pierce BCA assay (Thermo Fisher catalog # 23225) and equal protein content was used for Western blot. An equivalent volume of 2x Laemmli sample buffer (BioRad catalog # 1610737) containing freshly added 10% BME (M3148-25ML) was added to the worm samples prior to 10 min 95°C boiling. The samples were centrifuged (16,000 x g) and supernatants were collected. Samples were run on 4–15% tris-glycine mini gels (BioRad Mini-PROTEAN TGX stain-free gel catalog # 4568084) before transfer to PVDF membranes (BioRad catalog #1620177) at 4°C overnight at 0.3A in transfer buffer containing 25 mM Tris, 192 mM glycine, 20% methanol in milliQ water. Transferred membranes were blocked in 5% non-fat milk in Tris-buffered Saline (TBS) containing 0.1% Tween-20 (TBST) for 1 h. Primary antibodies in 3% non-fat milk in TBST were used for membranes incubation overnight at 4°C. The next day, membranes were washed 3 times in TBST and then incubated with secondary antibodies conjugated to IRdye 680LT (Licor, 1:20,000) in 3% non-fat milk in TBST at RT for 1 h, washed 3 times in TBST, and then imaged by BioRad ChemiDoc MP imaging system.

Primary antibodies used in this study: TUBA1 (mouse, Sigma-Aldrich catalog #T5168, 1:5000), GAPDH (mouse, EnCor Biotechnology catalog #MCA-1D4, RRID: AB_2107599, 1:5000).

Secondary antibody: IRDye 680LT Goat anti-Mouse IgG (LI-COR Biosciences catalog# 926-68020, RRID: AB_10706161).

### Statistics

The distributions of quantified data were tested for normality using Shapiro-Wilk test and appropriate statistical tests were carried out with GraphPad Prism. Error bars were shown as standard deviation unless otherwise indicated. Each figure legend lists the statistical test and the sample size. *ns*: p>0.05, *: p<0.05, **: p<0.005, ***: p<0.0005, and ****: p<0.0001. Exact p values were shown if p<0.05.

---

**Figure S1.**
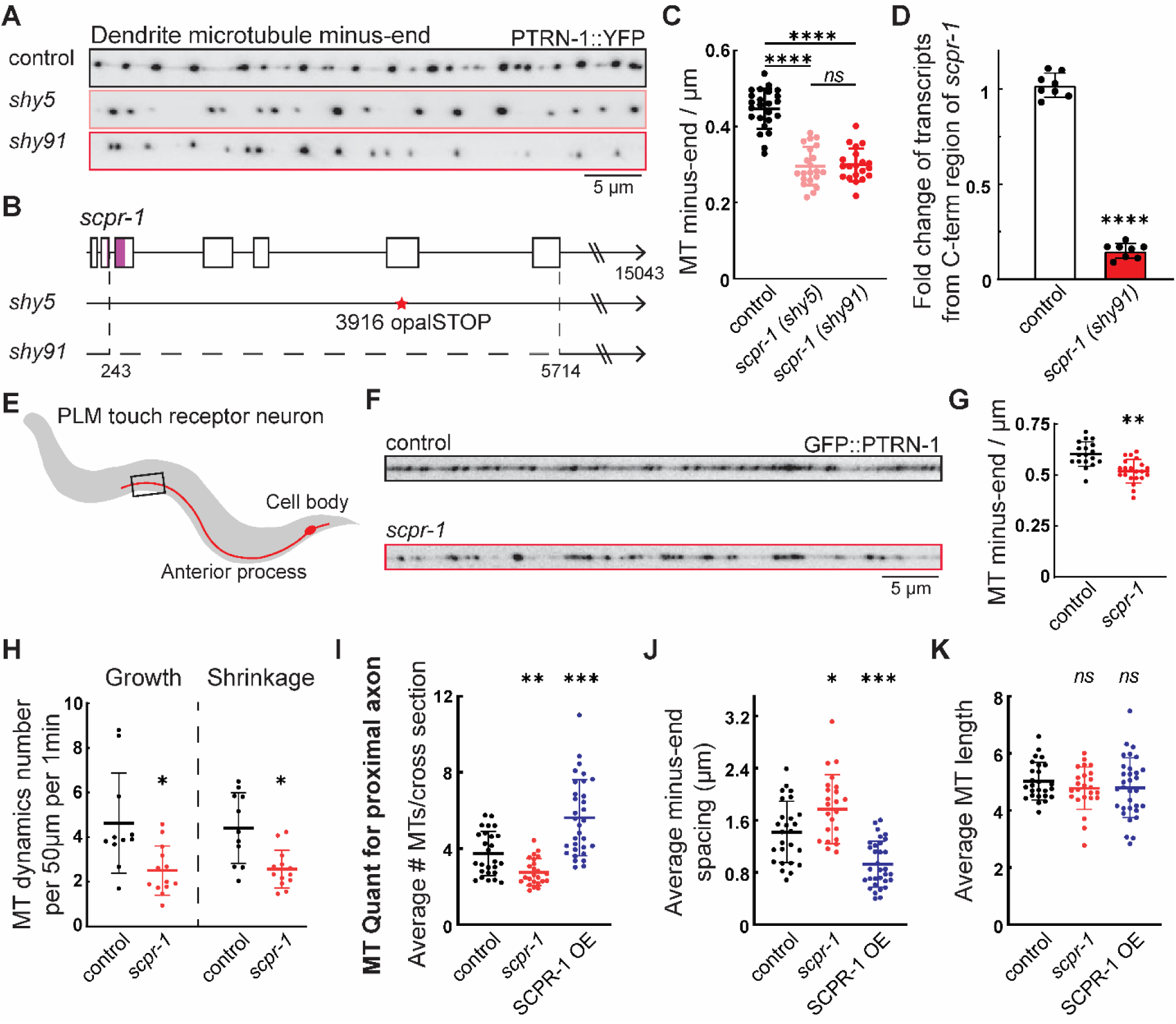
*scpr-1* mutants reduce MT content in *C. elegans* neurons. (A) Representative images of dendrite MT minus-end visualized by YFP-tagged PTRN-1 in control, *scpr-1 (shy5)*, and *scpr-1 (shy91)*. Scale bar = 5 μm. (B) Schematic of genetic map of N-terminus region of *scpr-1* gene with each *scpr-1* mutant. Magenta coated region indicates the CDS of conserved N-terminus SCAPER domain. Red star indicates the opalSTOP in *shy5* mutant, and dashed line indicates the deleted region in *shy91* mutant. (C) Quantification of MT minus-end number per μm in DA9 dendrite in control (N=20), *scpr-1 (shy5)* (N=21), and *scpr-1 (shy91)* (N=20). Brown-Forsythe and Welch ANOVA tests were used. ****: p<0.0001. *ns*: not significant. (D) Relative level of transcripts from C-terminus region of *scpr-1* from *scpr-1 (shy91)* vs N2 control by qRT-PCR. N=8 for both. Welch’s t test was used. ****: p<0.0001. (E) Schematic of C. elegans touch receptor neuron PLM. (F) Representative images of PLM anterior process MT minus-end visualized by GFP-tagged PTRN-1 in control and *scpr-1 (shy91)*. Scale bar = 5 μm. (G) Quantification of MT minus-end number per μm PLM anterior process in control (N=18) and *scpr-1 (shy91)* (N=21). Kolmogorov-Smirnov test was used. **: p=0.0017. (H) Quantification of MT dynamics event number DA9 dendrite in control and *scpr-1.* Control growth and shrinkage: N=11. *scpr-1* growth and shrinkage: N=13. Kruskal-Wallis tests were used. Growth event number in control vs *scpr-1* *: p=0.0107. Shrinkage event number in control vs *scpr-1* *: p=0.0157. (I-K) MT-Quant measurements and simulations of average number of MT per cross section, average MT minus-end spacing, and average MT length in DA9 axon for control (N=26), *scpr-1* (N=24), and SCPR-1 OE (N=32). Brown-Forsythe and Welch ANOVA tests were used. * (SCPR-1 OE vs control in J): p=0.0228. **: p<0.005, ***: p<0.0005. *ns*: not significant.

**Figure S2.**
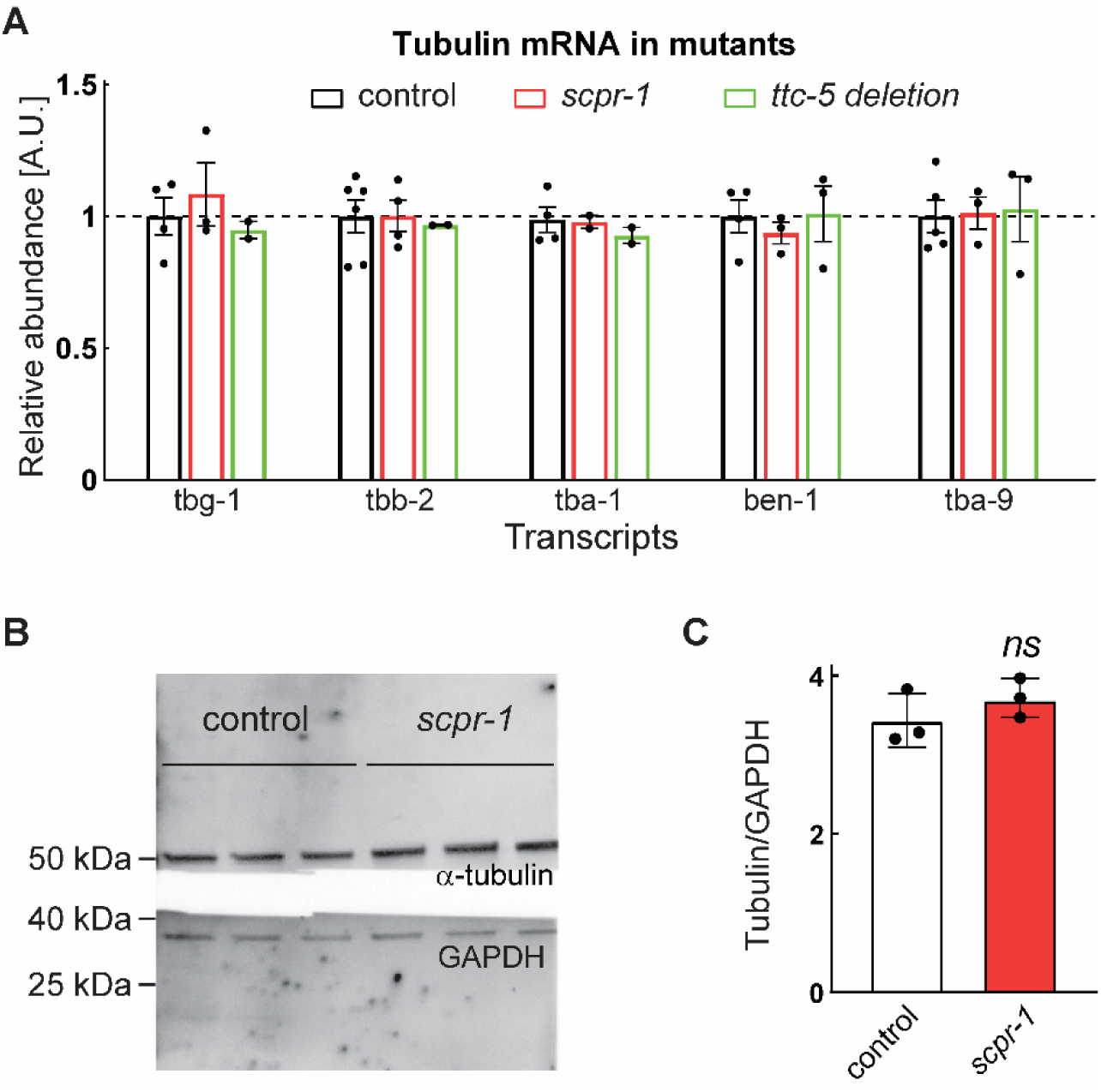
No change of tubulin mRNA or protein level in *scpr-1* mutant. (A) Relative mRNA level of γ-tubulin and MT subunits in control (N>=4), scpr-1 (N>=2), and ttc-5 deletion (N>=2). Welch’s t test was used. No significant difference compared to control. (B) Western blot of α-tubulin and GAPDH in control and *scpr-1* mutant. Three biological replicates for each were shown. (C) Quantification of relative α-tubulin protein level versus GAPDH in control and *scpr-1* mutant. N=3. Welch’s t test was used. *ns*: not significant.

**Figure S3.**
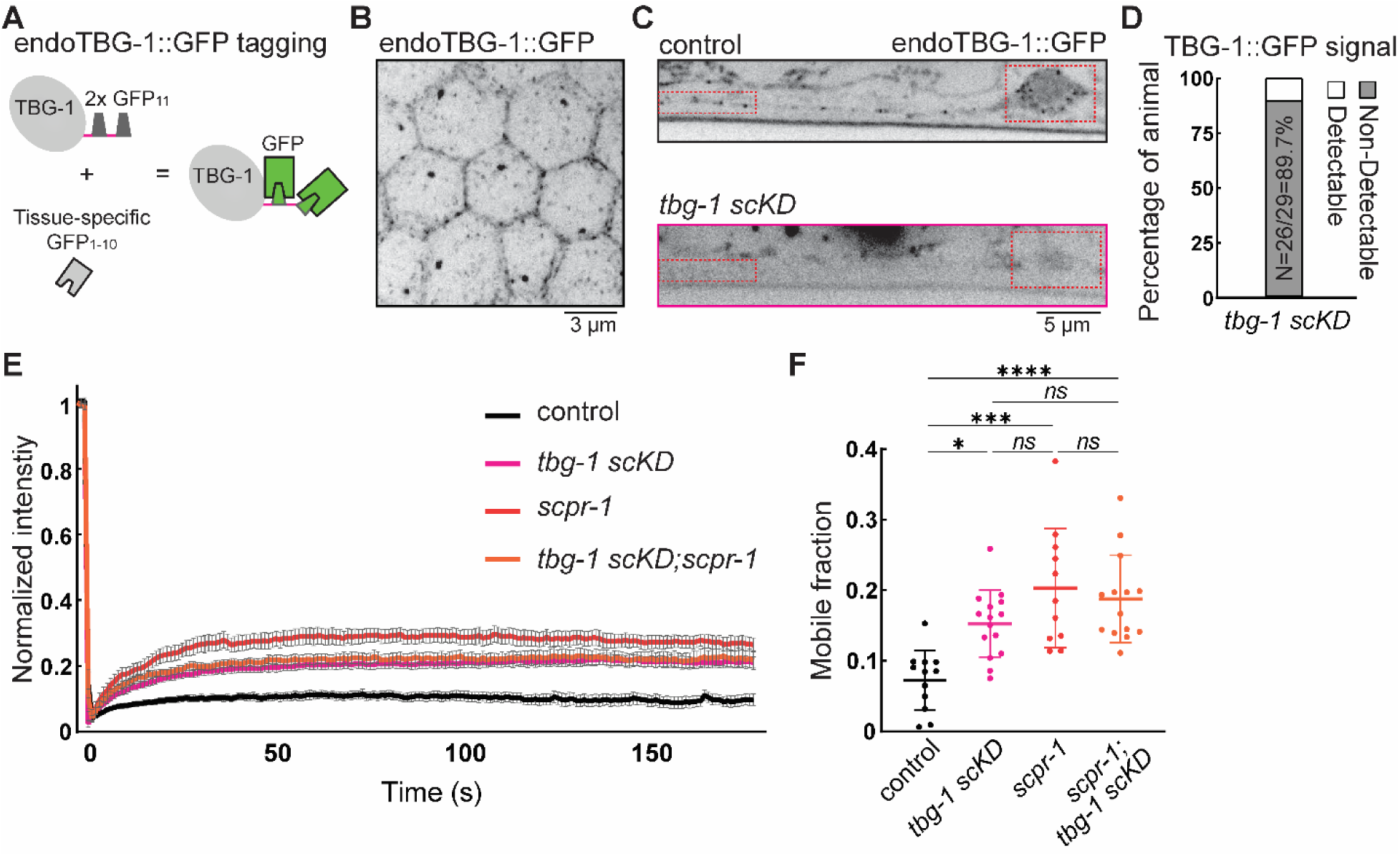
Endogenous γ-tubulin labeling and tubulin FRAP. (A) Schematic of γ-tubulin endogenous labeling for visualization. (B) Representative image of endogenous γ-tubulin signal in *C. elegans* germ cells. Scale bar = 3 μm. (C) Representative image of endogenous γ-tubulin signal in DA9 neuron with or without single cell knocking-down. Red rectangles highlight cell body and dendrite tip regions. Scale bar = 5 μm. (D) Quantification of percentage of animal that has detectable and non-detectable γ-tubulin signal in *tbg-1 scKD* condition. N=26/29 non-detectable. (E) Normalized intensity of GFP::TBA-1 FRAP in control (N=12), *tbg-1 scKD* (n=15), *scpr-1* (N=11), and *scpr-1*, *tbg-1 scKD; scpr-1* double mutant (N=14). Error bars were showed as S.E.M. (F) Calculation of mobile fraction of free tubulin in control (N=12), *tbg-1 scKD* (n=15), *scpr-1* (N=11), and *scpr-1*, *tbg-1 scKD; scpr-1* double mutant (N=14). *: p= 0.104 for *tbg-1 scKD* versus control, ***: p<0.0005, ****: p<0.0001 and *ns*: not significant by Kruskal-Wallis tests.

**Figure S4.**
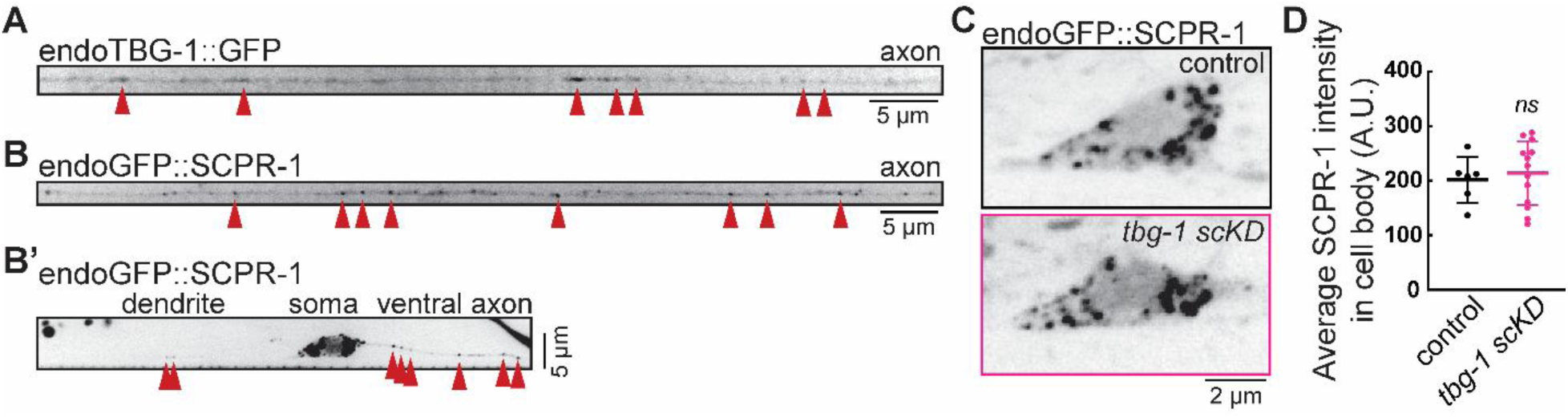
γ-tubulin and SCPR-1 are detected at low levels along axon and dendrite. (A) Representative image of endogenous γ-tubulin signal in DA9 axon. Scale bar = 5 μm. Punctate structures were highlighted by red arrow heads. (B and B’) Representative image of endogenous SCPR-1 signal in DA9 axon and dendrite, respectively. Punctate structures were highlighted by red arrow heads. Scale bar = 5 μm. (C) Representative image of endogenous SCPR-1 signal in DA9 cell body from control and *tbg-1 sckD*. (D) Quantification of average intensity of endogenous SCPR-1 signal in cell body from control (N=6) and *tbg-1 sckD* (N=13). *ns*: not significant by Mann-Whitney test.

**Figure S5.**
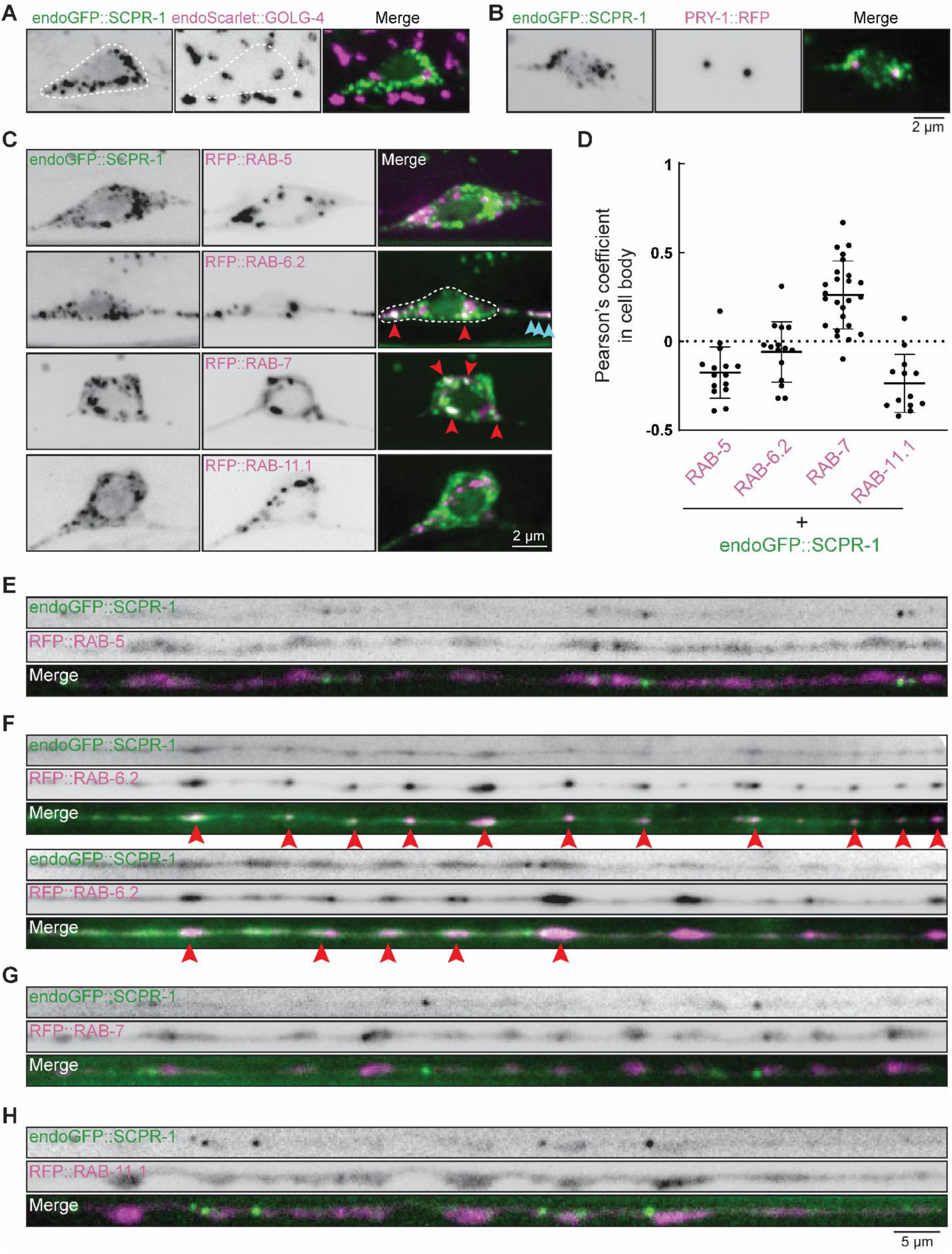
SCPR-1 localizes to endosomal structures. (A-C) Representative images of endogenous SCPR-1 with cellular structure markers in cell body. Endogenous tagged GOLGIN-4 (A), RFP tagged PRY-1 (B) and RAB-5, RAB-6.2, RAB-7, and RAB-11.1. SCPR-1 and markers were pseudo-colored green and magenta, respectively. White dashed lines highlighted cell body regions. Red arrow heads indicated colocalization in the cell body, and blue arrow heads indicated colocalization in the ventral axon. Scale bar = 2 μm. (D) Plot of Pearson’s coefficient of endogenous SCPR-1 with RFP tagged endosomal makers from (C). RAB-5 (N=16), RAB-6.2 (N=15), RAB-7 (N=25), and RAB-11.1 (N=13). (E) Representative images of endogenous SCPR-1 with RFP tagged endosomal markers in proximal axon. Two examples were shown from RAB-6.2. Red arrow heads highlighted colocalization. Scale bar = 5 μm.

**Figure S6.**
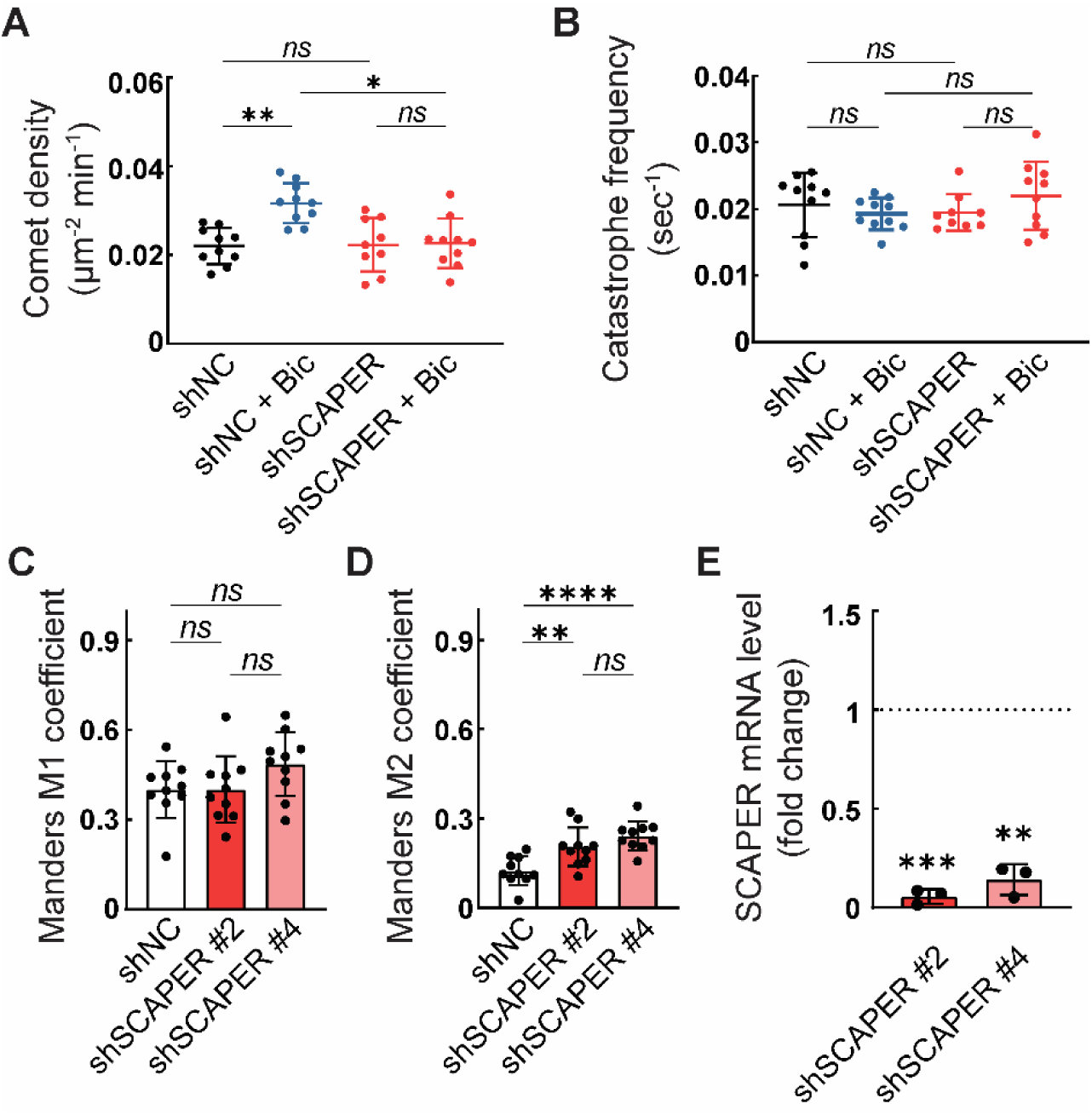
Mammalian SCAPER is required for γ-tubulin distribution and activity-dependent MT nucleation. (A and B) Quantification of EB3 comets density (A) and catastrophe frequency (B) in shRNA negative control (shNC) and SCAPER shRNA (shSCAPER) in control and Bic treatments (N>=9). * (shNC + Bic vs shSCAPER + Bic in A): p=0.0147, **: p<0.005, and *ns*: not significant by Kruskal-Wallis tests. (C and D) Manders coefficient of γ-tubulin signal to Synapsin 1/2 (C) or *vice versa* (D). n=10 field of view collected in N=2 independent experiments. **: p<0.005, ****: p<0.0001 by Mann-Whitney tests. (E) SCAPER mRNA levels measured as fold change of the non-coding control shNC (normalized to 1 as indicated by dashed line) after delivery of two different shRNA targeting SCAPER (shSCAPER#2 and #4) for 7 days. **: p<0.005, and ***: p<0.0005 by One sample t test.

**Figure S7.**
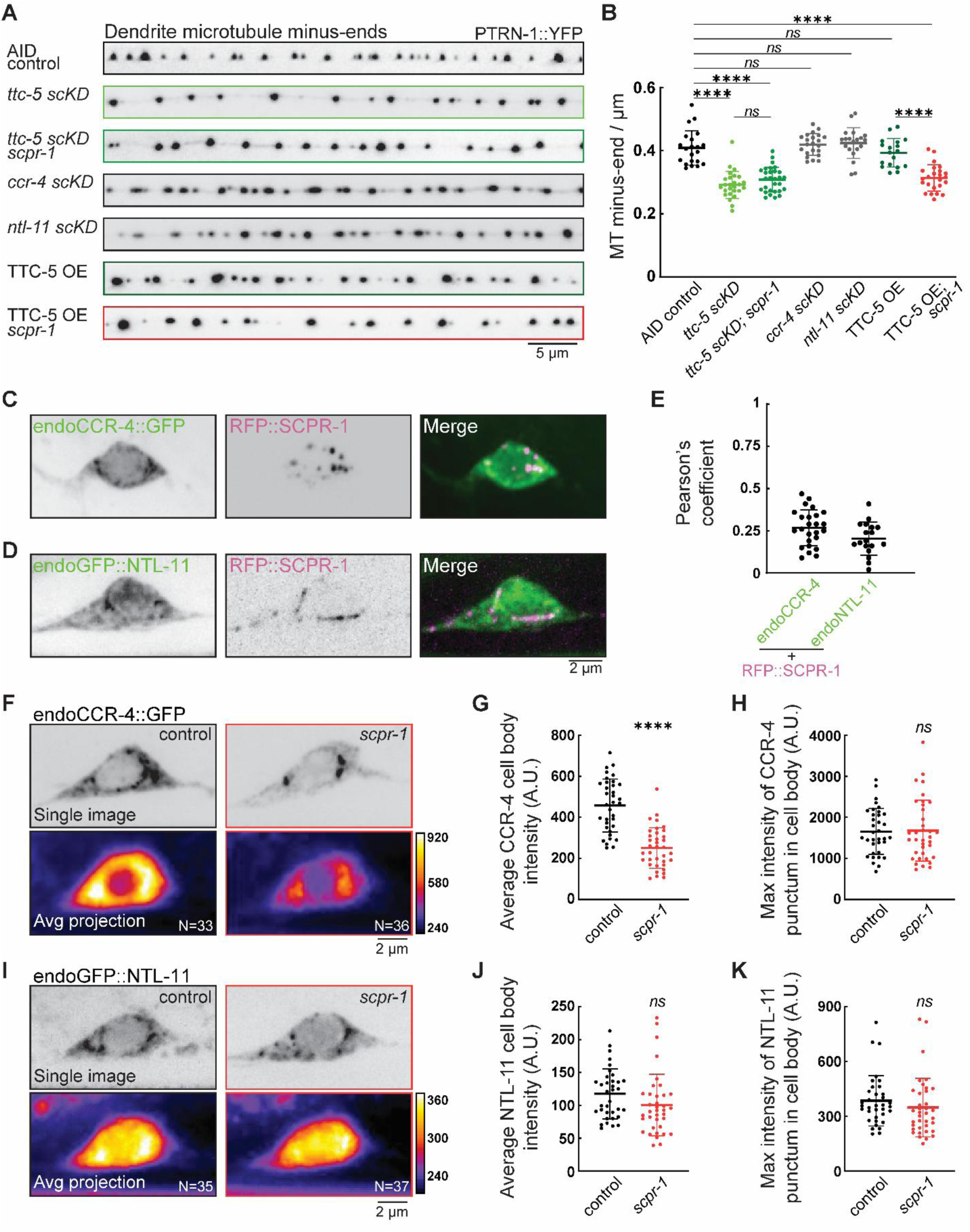
Localization and degradation phenotypes of CCR-4 and NTL-11. (A) Representative images of dendrite MT minus-end labeled by PTRN-1::YFP in AID control, *ttc-5 scKD, ttc-5 scKD; scpr-1* double mutant*, ccr-4 scKD, ntl-11 scKD,* TTC-5 overexpression, and TTC-5 overexpression in *scpr-1* mutant. Scale bar = 5 μm. (B) Quantification of MT minus-end number density in DA9 dendrite in AID control (N=21), *ttc-5 scKD* (N=26), *ttc-5 scKD; scpr-1* double mutant (N=29)*, ccr-4 scKD* (N=22), *ntl-11 scKD* (N=20), TTC-5 OE (N=18), and TTC-5 OE in *scpr-1* mutant (N=24). ****: p<0.0001, and *ns*: not significant by Brown-Forsythe and Welch ANOVA tests. (C and D) Representative cell body images of RFP::SCPR-1 transgene expressed in the endogenous CCR-4 (C) and NTL-11 (D) animals. Endogenous TTC-5::GFP and RFP::SCPR-1 were pseudo-colored as green and magenta, respectively. Scale bar = 2 μm. (E) Plot of Pearson’s coefficient from RFP::SCPR-1 with endogenous CCR-4::GFP (N=25) and GFP::NTL-11 (N=18) from cell body. (F and I) Representative images of endogenous CCR-4 (F) and NTL-11 (I) signal from control and *scpr-1* in cell body. Single images and average projection of multiple images pseudo-colored as heat maps for cell body were shown for control (N=33 and 35, respectively) and *scpr-1* (N=36 and 37, respectively). Spectrum bars shown as intensity [A. U.]. Scale bars = 2 μm. (G, H, J, K) Plot of average intensity of endogenous CCR-4 (G) and NTL-11 (J) and maximum intensity of CCR-4 (H) and NTL-11 (K) punctum, respectively, in cell body from control (N=35) and *scpr-1* (N=36 and 37, respectively). ****: p<0.0001, and *ns*: not significant by Mann-Whitney test.

**Figure S8.**
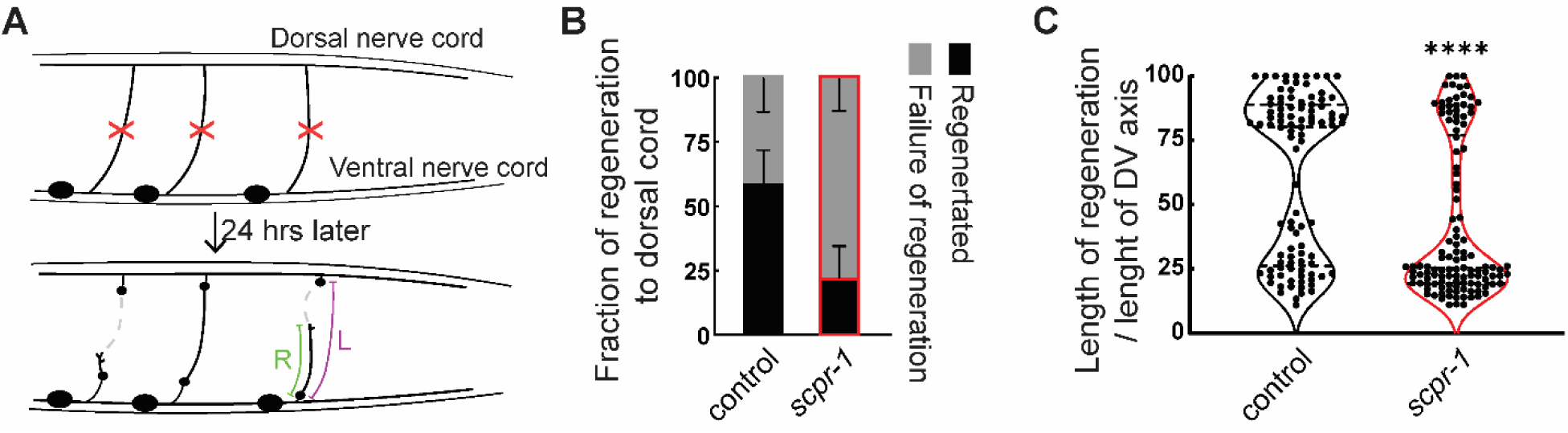
*scpr-1* is required for GABA neuron regeneration. (A) Schematic of axotomy in GABA neurons. Succeed regeneration was determined by regrowth of axon to the dorsal nerve cord. Relative regeneration length was calculated by R/L 24 hours after axotomy. (B) Plot of successful regeneration fraction from control (N=62/97) and *scpr-1* (N=39/118). (C) Plot of relative regeneration length / dorsal-ventral axis length in control (N=112) and *scpr-1* (N=134). Kolmogorov-Smirnov test was applied. ****: p<0.0001.

**Table S1.** C. elegans strains generated and used in this study.

| Strain ID | Genotype | Source |
| --- | --- | --- |
| TV21554 | wyls802[itr-1p::ptrn-1::yfp, odr-1p::gfp] | Shen lab |
| MTS536 | *scpr-1(shy5)*; wyls802 | this study |
| MTS1344 | *scpr-1(shy91)*; wyls802 | this study |
| MTS1628 | *scpr-1(shy91)*; wyls802; shyEx420[mig-13p::rfp::scpr-1] | this study |
| MTS2858 | wyls802; shyEx420[mig-13p::rfp::scpr-1] | this study |
| MTS2922 | *scpr-1(shy91)*; wyls802, wySi265[unc-4cp::CRE]; shyEx702[mig-13p::stop::rfp::hs_scaper] | this study |
| MTS226 | *tba-1(ok1135)*; wySi265; wyls463[mig-13p::stop::gfp::tba-1] | this study |
| MTS1490 | *tba-1(ok1135)*, *scpr-1(shy91)*; wySi265; wyls463[mig-13p::stop::gfp::tba-1] | this study |
| MTS2848 | *tba-1(ok1135)*, *scpr-1(shy91)*; wySi265; wyls463; shyEx553[mig-13p::rfp::scpr-1] | this study |
| MTS1576 | *tba-1(ok1135)*; wySi265; wyls463; shyEx413[mig-13p::stop::rfp::delCC_ptrn-1] | this study |
| MTS1660 | *tba-1(ok1135)*, *scpr-1(shy91)*; wySi265; wyls463; shyEx413 | this study |
| MTS1845 | *tba-1(ok1135)*; wySi265; wyls463; shyEx451[mig-13p::stop::rfp::delCC_ptrn-1, mig-13p::scpr-1] | this study |
| MTS2645 | wySi265; *tbg-1(shy152[tbg-1::2xgfp11])*; shyls123[mig-13p::stop::gfp1-10] | this study |
| MTS2758 | wyls802, wySi265; *tbg-1(shy174[tbg-1::1xgfp11::zf1])*, shyls111[mig-13p::stop::zif-1] | this study |
| MTS2759 | *scpr-1(shy91)*; wyls802, wySi265; *tbg-1(shy174)*, shyls111 | this study |
| MTS2856 | *scpr-1(shy91)*; wyls802, wySi265; *tbg-1(shy174)*, shyls111; shyEx420 | this study |
| MTS2855 | wyls802; wySi265; *tbg-1(shy174)*, shyls111; shyEx420 | this study |
| MTS2794 | *tba-1(ok1135)*; wySi265; *tbg-1(shy174)*, shyls111; wyls463 | this study |
| MTS2798 | *tba-1(ok1135)*, *scpr-1(shy91)*; wySi265; *tbg-1(shy174)*, shyls111; wyls463 | this study |
| MTS2847 | *tba-1(ok1135)*, *scpr-1(shy91)*; wySi265; *tbg-1(shy174)*, shyls111; wyls463; shyEx553 | this study |
| MTS2857 | *tbg-1(shy174)*, shyls111; shyls35[mig-13p::gfp1-10] | this study |
| MTS2771 | *tbg-1(shy174)*, *shyls111*; *wySi265*; *shyls35* | this study |
| MTS2851 | *scpr-1(shy113[7xgfp11::scpr-1])*; *wySi265*; *shyls123* | this study |
| MTS2845 | *scpr-1(shy91)*; *wySi265*; *tbg-1(shy152)*; *shyls123*; *shyEx553* | this study |
| MTS2844 | *wySi265*; *tbg-1(shy152)*; *shyls123*; *shyEx553* | this study |
| MTS2904 | *scpr-1(shy113)*; *wySi265*; *tbg-1(shy174)*, *shyls111*; *shyls35* | this study |
| PHX6547 | *golg-4(syb6547[wormScarlet::golg-4])* | CGC/Hobert lab <sup>89</sup> |
| MTS3470 | *golg-4(syb6547[wormScarlet::golg-4])*; *scpr-1(shy113)*; *shyls35* | this study |
| MTS3075 | *scpr-1(shy113)*; *wySi265*; *shyls129[mig-13p::stop::gfp1-10]*; *shyEx770[mig-13p::stop::pry-1::rfp]* | this study |
| MTS2888 | *scpr-1(shy113)*; *wySi265*; *shyls129*; *shyEx678[mig-13p::rfp::rab-5]* | this study |
| MTS3078 | *scpr-1(shy113)*; *wySi265*; *shyls129*; *shyEx773[mig-13p::stop::rfp::rab-6.2]* | this study |
| MTS2936 | *scpr-1(shy113)*; *wySi265*; *shyls129*; *shyEx713[mig-13p::stop::rfp::rab-7]* | this study |
| MTS2939 | *scpr-1(shy113)*; *wySi265*; *shyls129*; *shyEx716[mig-13p::stop::rfp::rab-11.1]* | this study |
| MTS2842 | *ebp-2(shy70[ebp-2::7xgfp11])*; *shyls32[mig-13p::gfp1-10]*; *shyEx553* | this study |
| MTS2843 | *scpr-1(shy91)*; *ebp-2(shy70)*; *shyls32*; *shyEx553* | this study |
| MTS3108 | *shyEx553*; *ebp-2(shy70)*, *wySi265*; *tbg-1(shy174)*, *shyls111*; *shyls32* | this study |
| MTS3248 | *ttc-5(shy254[ttc-5 deletion])*; *wyls802* | this study |
| MTS3249 | *ttc-5(shy256[ttc-5 r138a])*; *wyls802* | this study |
| MTS3322 | *ttc-5(shy254)*, *scpr-1(shy91)*; *wyls802* | this study |
| MTS3504 | *ttc-5(shy254)*; *wyls802*; *shyEx420* | this study |
| MTS3106 | *ttc-5(shy225[ttc-5::AID::4xgfp11])*; *shyls35*; *shyEx553* | this study |
| MTS3297 | *ttc-5(shy254)*, *scpr-1(shy113)*; *shyls35* | this study |
| MTS3290 | *ttc-5(shy225)*, *scpr-1(shy91)*; *shyls35* | this study |
| MTS3400 | *ttc-5(shy254)*; *tbg-1(shy152)*; *shyls35* | this study |
| MTS3401 | *ttc-5(shy256)*; *tbg-1(shy152)*; *shyls35* | this study |
| MTS3172 | *ttc-5(shy225)*; *wyls802*; *shyEx810[mig-13p::TIR1]* | this study |
| MTS3190 | *ttc-5(shy225), scpr-1(shy91)*; wyls802; shyEx810 | this study |
| MTS3197 | wyls802,wySi265; shyEx693[mig-13::stop::ttc-5::rfp] | this study |
| MTS3198 | *scpr-1(shy91)*; wyls802,wySi265; shyEx693 | this study |
| MTS3255 | shyEx553; *ccr-4(shy231[ccr-4::AID::4xgfp11])*; shyls36[mig-13p::gfp1-10] | this study |
| MTS3256 | shyEx553; *ntl-11(shy240[AID::4xgfp11::ntl-11])*; shyls36 | this study |
| MTS3199 | *ccr-4(shy231)*; shyls36 | this study |
| MTS3200 | *ntl-11(shy240)*; shyls36 | this study |
| MTS3293 | *scpr-1(shy91)*; *ccr-4(shy231)*; shyls36 | this study |
| MTS3298 | *scpr-1(shy91)*; *ntl-11(shy240)*; shyls36 | this study |
| DC19 | *bus-5(br19)* | CGC |
| MTS3275 | *scpr-1(shy91)*; *bus-5(br19)* | this study |
| MTS3294 | *ttc-5(shy254)*; *bus-5(br19)* | this study |
| MTS2804 | wySi265; *tbg-1(shy174)*, shyls111; shyEx654[itr-1p::myr::rfp] | this study |
| MTS2805 | *scpr-1(shy91)*; *tbg-1(shy174)*, shyls111; wySi265; shyEx655[itr-1p::myr::rfp] | this study |
| MTS2008 | *scpr-1(shy113)*; shyls35; shyEx493[itr-1p::myr::rfp] | this study |
| MTS2120 | *tbg-1(shy152)*; shyls35; shyEx493[itr-1p::myr::rfp] | this study |
| MTS2288 | *scpr-1(shy91)*; *tbg-1(shy152)*; shyls35; shyEx546[itr-1p::myr::rfp] | this study |
| XE2082 | wpls109[itr-1p::mCherry] | Hammarlund lab |
| MTS2233 | *scpr-1(shy91)*; wpls109 | this study |
| MTS2804 | wySi265; *tbg-1(shy174)*, shyls111; shyEx654[itr-1p::myr::rfp] | this study |
| MTS2805 | *scpr-1(shy91)*; *tbg-1(shy174)*, shyls111; wySi265; shyEx655[itr-1p::myr::rfp] | this study |
| MTS2008 | *scpr-1(shy113)*; shyls35; shyEx493[itr-1p::myr::rfp] | this study |
| MTS2120 | *tbg-1(shy152)*; shyls35; shyEx493[itr-1p::myr::rfp] | this study |
| MTS2288 | *scpr-1(shy91)*; *tbg-1(shy152)*; shyls35; shyEx546[itr-1p::myr::rfp] | this study |
| XE1873 | oxls12[unc-47p::gfp] | Hammarlund lab |
| MTS1548 | *scpr-1(shy91)*; oxls12 | this study |

**Table S2.**
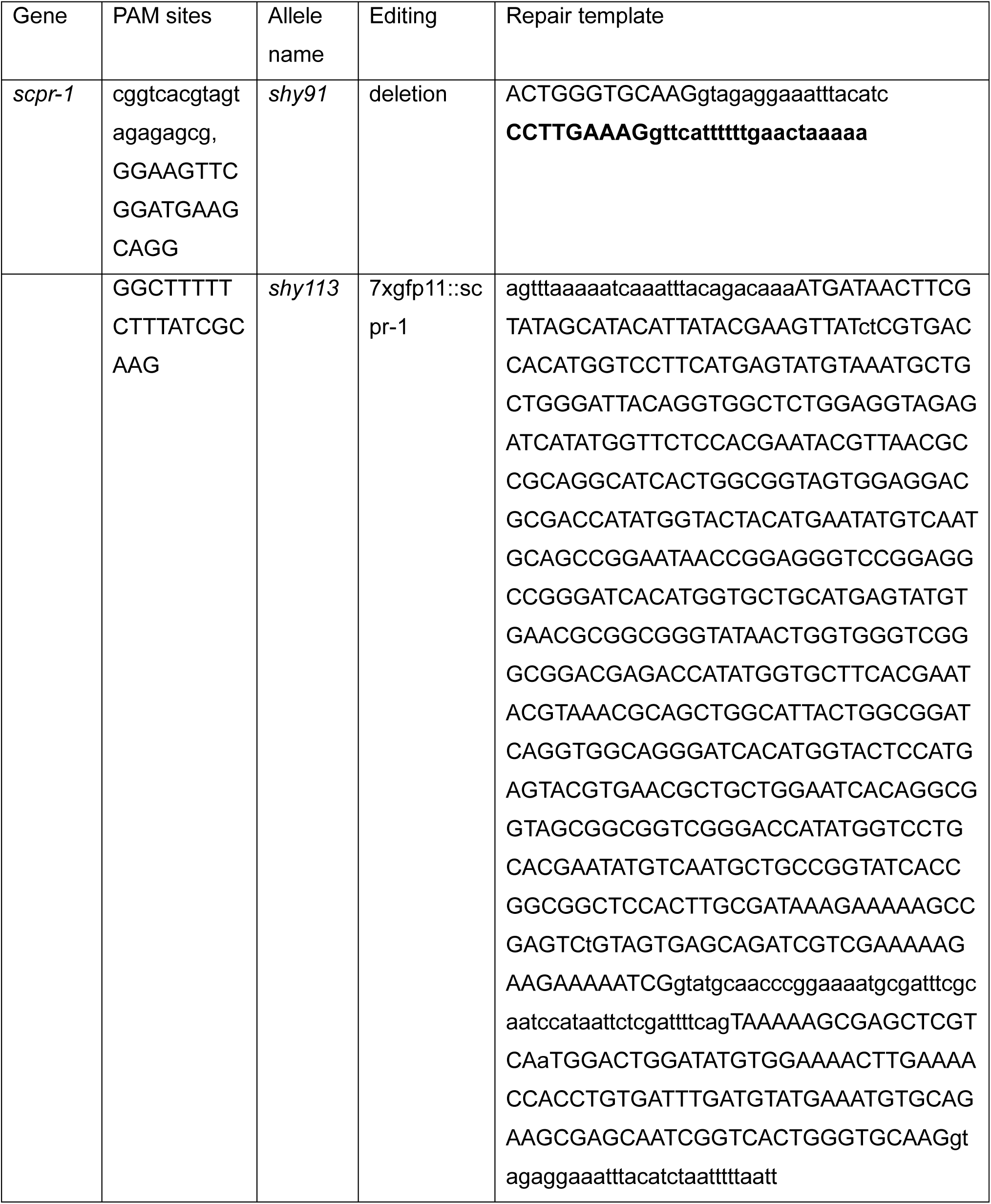
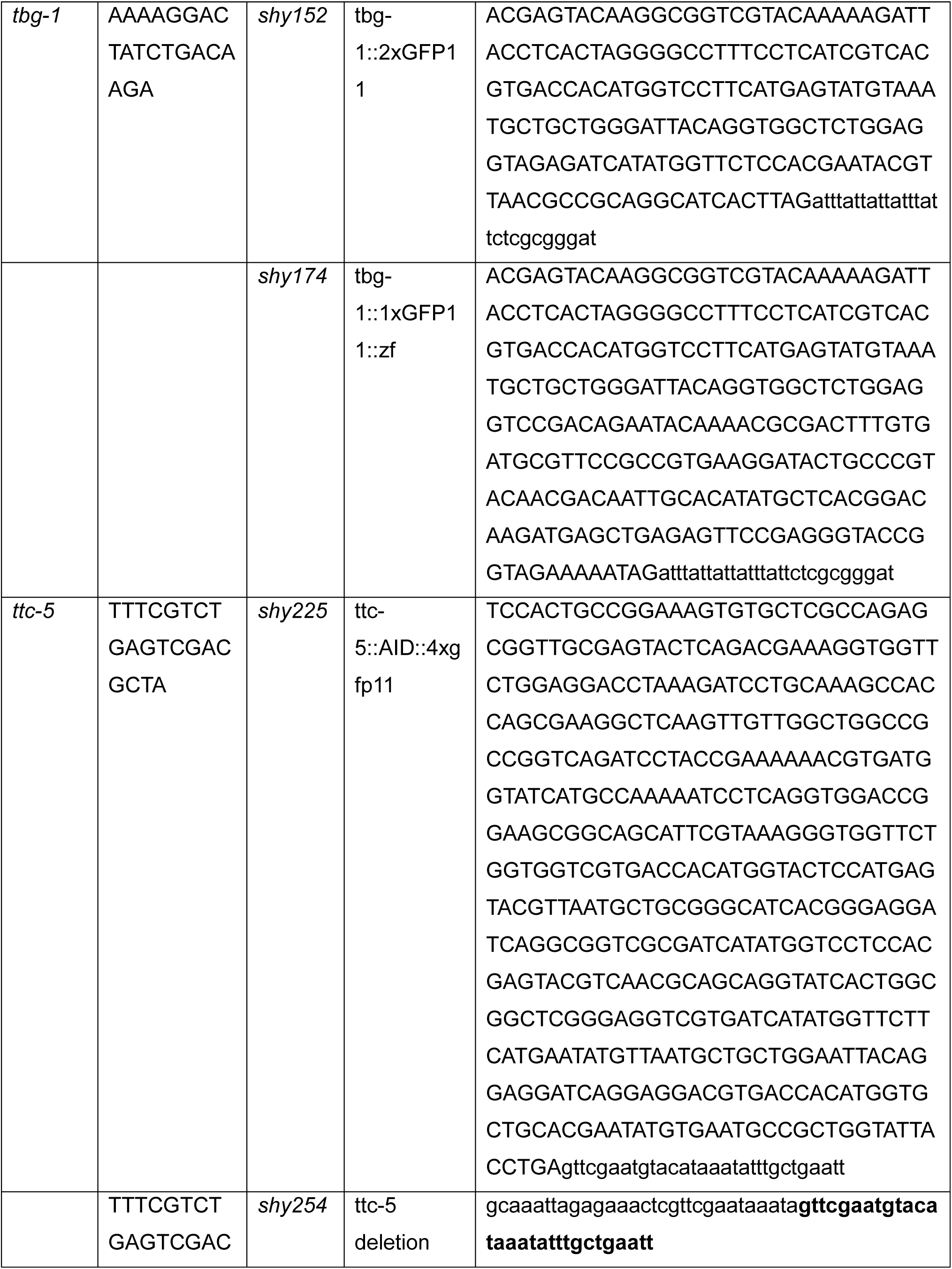
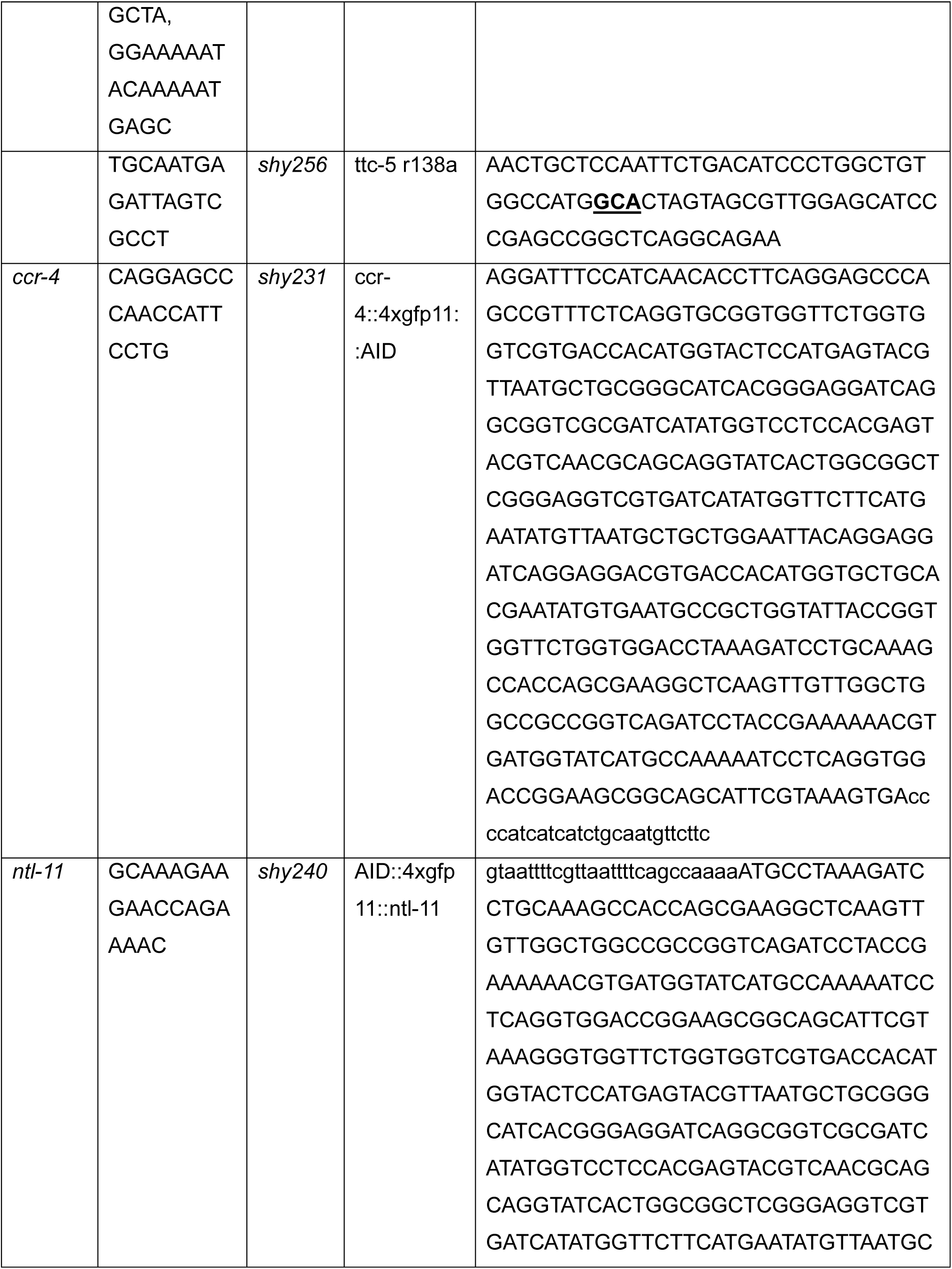
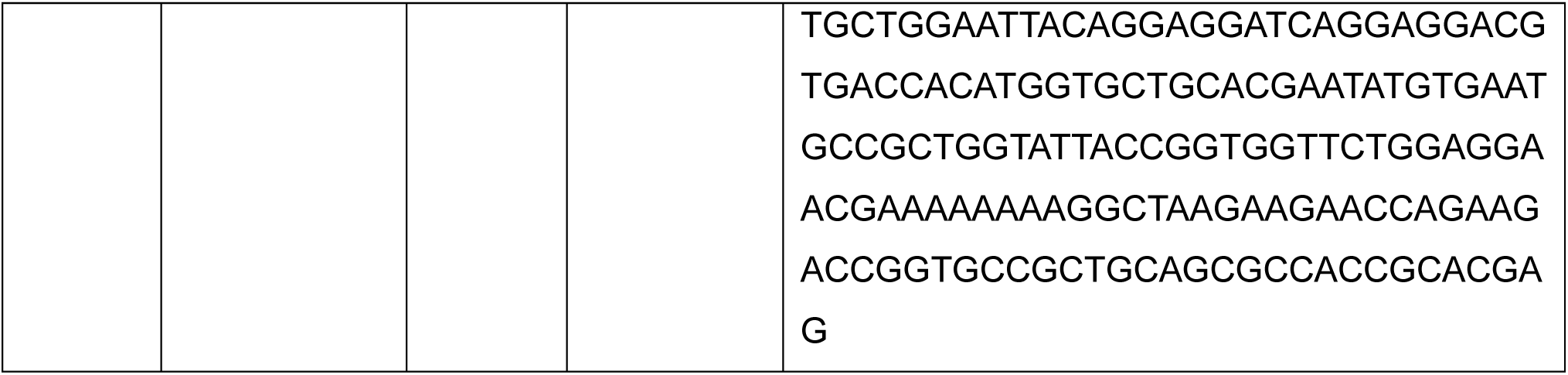
gRNA used in this study.

**Table S3.** Plasmids generated and used in this study.

| Identifier | Reagent | Arrays |
| --- | --- | --- |
| pYX15 | mig-13p::rfp::scpr-1 | shyEx420, shyEx553 |
| pYX16 | mig-13p::stop::rfp::delCC_ptrn-1 | shyEx413, shyEx451 |
| pYX69 | mig-13p::stop::rfp::hs_scaper | shyEx702 |
| pYX8 | mig-13p::scpr-1 | shyEx451 |
| pYX54 | mig-13p::stop::gfp1-10 | shyls123, shyls129 |
| pYX55 | mig-13p::stop::zif-1 | shyls111 |
| pYX84 | mig-13p::stop::pry-1::rfp | shyEx770 |
| pYX62 | mig-13p::rfp::rab-5 | shyEx678 |
| pYX86 | mig-13p::stop::rfp::rab-6.2 | shyEx773 |
| pYX75 | mig-13p::stop::rfp::rab-7 | shyEx713 |
| pYX76 | mig-13p::stop::rfp::rab-11.1 | shyEx716 |
| pOVG13 | Mig-13p::tir1 | shyEx810 |
| pYX67 | mig-13p::stop::ttc-5::rfp | shyEx693 |
| pCF6 | itr-1p::myr::rfp | shyEx493, shyEx546, shyEx654, shyEx655 |
| pRF4 | rol-6(su1006) |  |

**Table S4.**
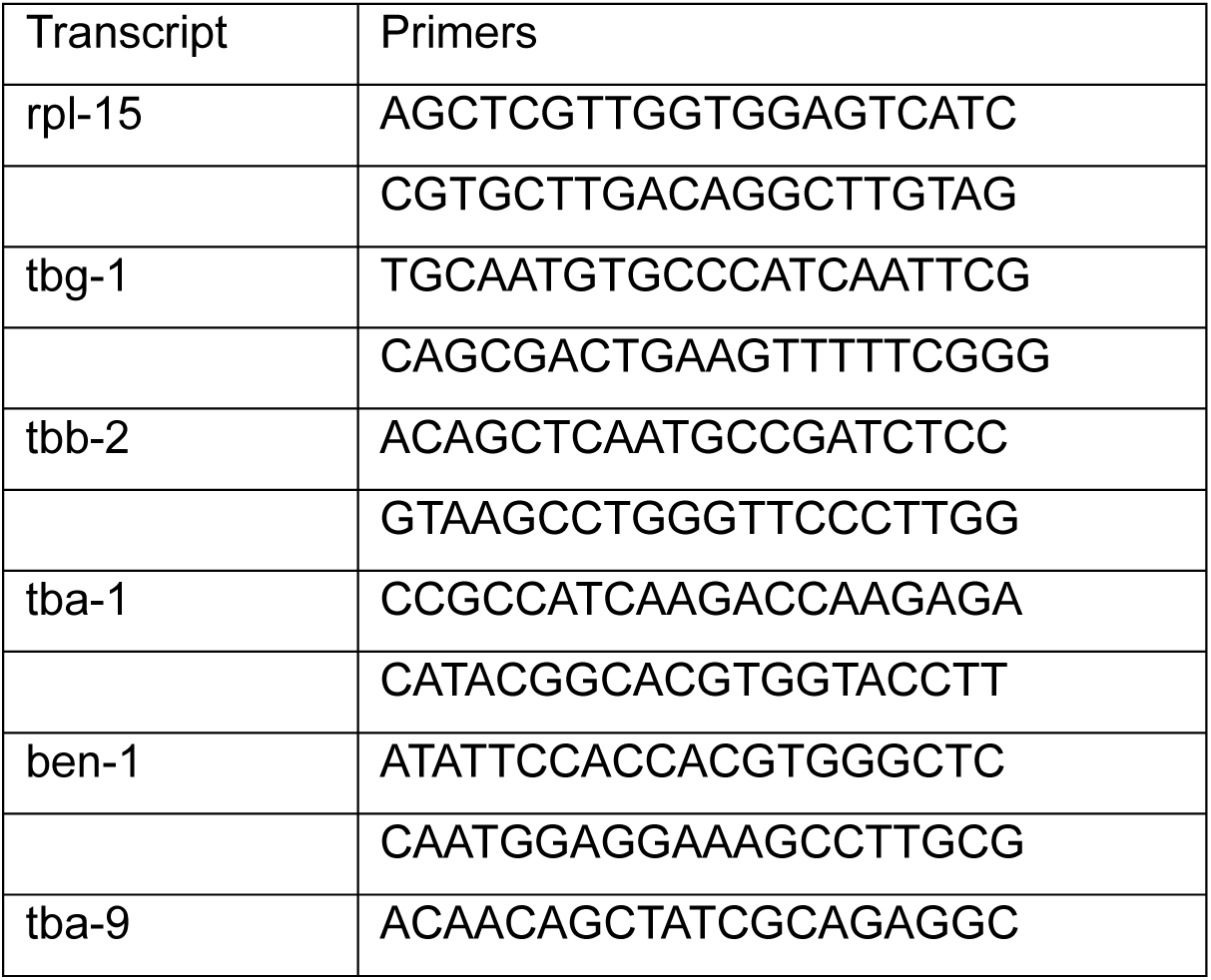
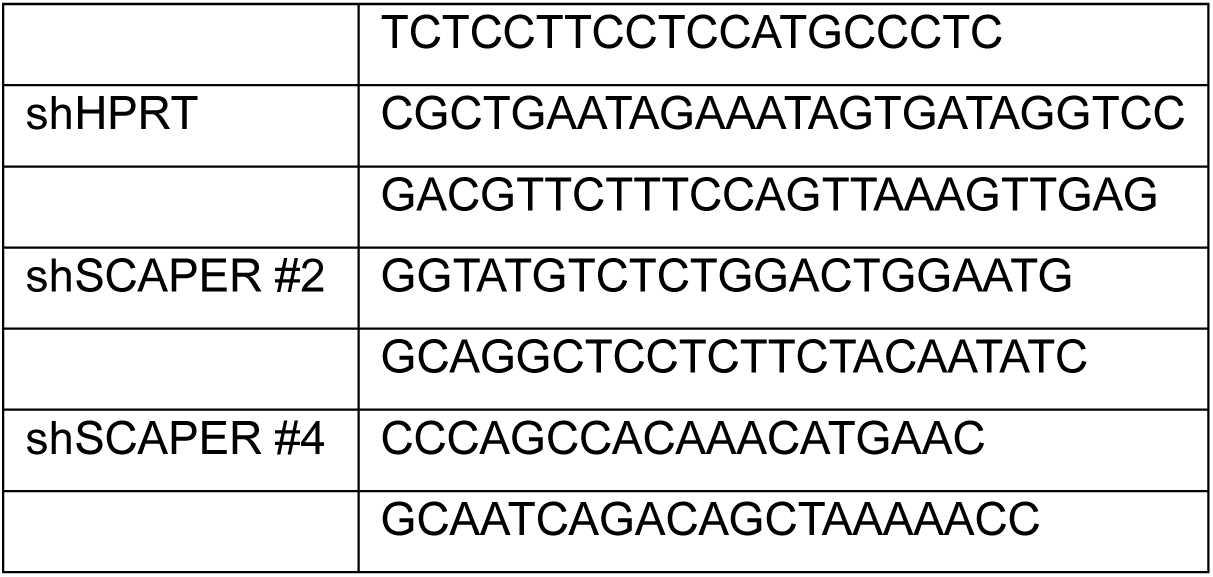
Primers for qRT-PCR used in this study.

